# Two *rsaM* homologues encode central regulatory elements modulating quorum sensing expression in *Burkholderia thailandensis*

**DOI:** 10.1101/192625

**Authors:** Servane Le Guillouzer, Marie-Christine Groleau, Eric Déziel

## Abstract

The bacterium *Burkholderia thailandensis* possesses three conserved *N*-acyl-_L_-homoserine lactone (AHL) quorum sensing (QS) systems designated BtaI1/BtaR1 (QS-1), BtaI2/BtaR2 (QS-2), and BtaI3/BtaR3 (QS-3). These QS-systems are associated with the biosynthesis of *N*-octanoyl-homoserine lactone (C_8_-HSL), *N*-3-hydroxy-decanoyl-homoserine lactone (3OHC10-HSL), as well as *N*-3-hydroxy-octanoyl-homoserine lactone (3OHC_8_-HSL), which are produced by the LuxI-type synthase BtaI1, BtaI2, and BtaI3, and modulated by the LuxR-type transcriptional regulators BtaR1, BtaR2, and BtaR3. Both *btaR1*/*btaI1* and *btaR2*/*btaI2* gene clusters contain an additional gene that is conserved in the *Burkholderia* genus, homologous to a gene coding for the negative AHL biosynthesis modulatory protein RsaM originally identified in the phytopathogen *Pseudomonas fuscovaginae*, and hence designated *rsaM1* and *rsaM2*. We have characterized the function of these two *rsaM* homologues and demonstrated their involvement in the regulation of AHLs biosynthesis in *B. thailandensis* strain E264. We measured the production of C_8_-HSL, 3OHC_10_-HSL, and 3OHC_8_-HSL by liquid chromatography coupled to tandem mass spectrometry (LC-MS/MS) in the wild-type strain and in the *rsaM*1^-^ and *rsaM*2^-^ mutants, and monitored the transcription of *btaI*1, *btaI*2, and *btaI*3 using chromosomal mini-CTX-*lux* transcriptional reporters. The expression of *btaR*1, *btaR*2, and *btaR*3 was also measured by quantitative everse-transcription PCR (qRT-PCR). We demonstrate that the QS-1 system is repressed by RsaM1, whereas RsaM2 principally represses the QS-2 system. We also found that both *rsaM1* and *rsaM2* are QS-controlled, as well as negatively auto-regulated. We conclude that RsaM1 and RsaM2 are an integral part of the QS modulatory circuitry of *B. thailandensis*, and play a major role in the hierarchical and homeostatic organization of the QS-1, QS-2, and QS-3 systems.

**Importance:** Quorum sensing (QS) is a global regulatory mechanism of genes expression depending on bacterial density. QS is commonly involved in the coordination of genes expression associated with the establishment of host-pathogen interactions and acclimatization to the environment. We present the functional characterization of the two *rsaM* homologues designated *rsaM1* and *rsaM2* in the regulation of the multiple QS systems coexisting in the non-pathogenic bacterium *Burkholderia thailandensis*, widely used as a model system for the study of the pathogen *Burkholderia pseudomallei*. We found that inactivation of these *rsaM* homologues, which are clustered with the other QS genes, profoundly affects the QS regulatory circuity of *B. thailandensis*. It is proposed that these genes code for QS repressors and we conclude that they constitute essential regulatory components of the QS modulatory network of *B. thailandensis*, and provide additional layers of regulation to modulate the expression of QS-controlled genes, including those encoding virulence/survival factors and linked to environmental adaptation in *B. pseudomallei*.

## Introduction

Quorum sensing (QS) is a widespread bacterial intercellular communication system that coordinates expression of specific genes in a cell density-dependent manner(1). QS is mediated by diffusible signaling molecules, called autoinducers, which are synthesized and secreted in response to fluctuations in cell density. They accumulate in the environment as bacterial growth progresses until a threshold concentration is reached allowing the coordination of the expression of specific genes. Gram-negative bacteria typically possess homologues of the LuxI/LuxR system initially characterized in the bioluminescent marine bacterium *Vibrio fischeri* (2). The signaling molecules *N*-acyl-L-homoserine lactones (AHLs) are produced by the LuxI-type synthases. These AHLs activate the LuxR-type transcriptional regulators that modulate the expression of QS target genes, which usually contain a *lux*-box sequence in their promoter region. These genes include a *luxI* homologue encoding the AHL synthase, resulting in a typical self-inducing loop of AHLs (3).

The *Burkholderia* genus encompasses heterogeneously species colonizing diverse ecological niches, such as soil, water, plants, as well as animals, including humans (4). The *Burkholderia cepacia* complex (Bcc), for instance comprises notable opportunistic human pathogens deleterious in both cystic fibrosis (CF) individuals and immunocompromised patients (5). Bcc members carry *luxI* and *luxR* homologues referred as *cepI* and *cepR* genes, respectively. These genes are constitutive of the AHL-based QS system designated CepI/CepR (6). The LuxI-type synthase CepI is responsible for *N*-octanoyl-homoserine lactone (C_8_-HSL) biosynthesis, which generally constitutes the predominant AHL found in the *Burkholderia* genus (6). The LuxR-type transcriptional regulator CepR modulates the expression of QS target genes in conjunction with C_8_-HSL, including the *cepI* gene creating the QS typical auto-regulation loop (6). The genetic organization of *cepI* and *cepR* is conserved among *Burkholderia* spp. (7). These genes are generally separated by a gene encoding an RsaM-like protein that was originally identified in the plant pathogen *Pseudomonas fuscovaginae* and shown to be a major negative regulator of both AHLs biosynthesis and expression of the AHL synthase-coding genes (8). It was also recently reported to act as a global regulator mediating the expression of numerous genes through and out of the QS regulon in *P. fuscovaginae* (9). The function of RsaM-like proteins has been investigated in the *Burkholderia* genus, and could be important for balancing and fine-tuning of QS-dependant regulation (10). RsaM-like proteins do not possess any sequence similarity with biochemically or structurally characterized proteins, such as DNA-binding motifs, and constitute single-domain proteins with unique topology presenting a novel fold (11). Their precise underlying regulatory mechanism is currently unknown.

The non-pathogenic tropical soil saprophyte *Burkholderia thailandensis*, as well as the closely-related human pathogen *Burkholderia pseudomallei*, both encode two conserved RsaM-like proteins of uncharacterized function (7). The genome of *B. thailandensis* contains three LuxI/LuxR type QS systems designated BtaI1/BtaR1 (QS-1), BtaI2/BtaR2 (QS-2), and BtaI3/BtaR3 (QS-3). These QS systems are also found in *B. pseudomallei*, and were reported to be involved in the regulation of several virulence genes and to be essential to its pathogenicity (12). The QS-1, QS-2, and QS-3 systems were recently reported to be hierarchically and homeostatically organized, and integrated into an intricate modulatory network, including transcriptional and post-transcriptional interactions between each QS circuits (13). The QS-1 AHL synthase BtaI1 produces C_8_-HSL (14) which associates with the BtaR1 transcriptional regulator to activate the expression of the *btaI*1 gene (13, 15). The QS-2 system is responsible for the biosynthesis of both *N*-3-hydroxy-decanoyl-homoserine lactone (3OHC_10_-HSL) and *N*-3-hydroxy-octanoyl-homoserine lactone (3OHC_8_-HSL) (16). The *btaI*2 gene, which encodes the BtaI2 synthase, is positively controlled by the BtaR2 transcriptional regulator in association with 3OHC_10_-HSL and 3OHC_8_-HSL (13, 16). The QS-3 system is composed of the BtaR3 transcriptional regulator and the BtaI3 synthase responsible for 3OHC_8_-HSL production as well (14). The *btaI*3 gene encoding BtaI3 is activated by BtaR3 (13). While both the QS-1 and QS-2 gene clusters include an *rsaM* homologue, no *rsaM* gene is present in the vicinity of *btaI*3/*btaR3* (7).

The central aim of this study was to further explore the QS modulatory circuitry of *B. thailandensis* E264 to identify additional transcriptional and/or post-transcriptional regulators of the QS-1, QS-2, and QS-3 systems. We functionally characterized the role of *rsaM1* and *rsaM2* in the regulation of the *B. thailandensis* E264 AHL-based QS circuits. We established that they negatively modulate AHLs biosynthesis and that they are finely integrated into the complex QS circuitry of *B. thailandensis* E264. This study provides new insights on the intricate interplay existing between the QS systems of *B. thailandensis*, and is essential in unraveling the regulatory mechanism underlying QS-dependent genes expression, including those encoding virulence/survival factors and linked to environmental adaptation in *B. pseudomallei*.

## Materials and methods

### Bacterial strains and culture conditions

The bacterial strains used in this study are listed in **Table S1**. Unless otherwise stated, all bacteria were cultured at 37°C in Tryptic Soy Broth (TSB; BD Difco(tm), Mississauga, ON, Canada), with shaking (240 rpm) in a TC-7 roller drum (New Brunswick, Canada), or in Petri dishes containing TSB solidified with 1.5% agar. When required, antibiotics were used at the following concentrations: 200 μg/mL tetracycline (Tc) and 100 μg/mL trimethoprim (Tp) for *B. thailandensis* E264, while Tc was used at 15 μg/mL for *Escherichia coli* DH5α. All measurements of the optical density (OD_600_) were acquired with a Thermo Fisher Scientific NanoDrop® ND-1000 Spectrophotometer.

### Plasmids construction

All plasmids used in this study are described in **Table S2**. Amplification of *btaR*2 was conducted from genomic DNA of *B. thailandensis* E264 using appropriate primers (**Table S3**). The amplified products were digested with the FastDigest restriction enzymes *Bam*HI and *Hind*III (Thermo Fisher Scientific) and ligated using T4 DNA ligase (Bio Basic, Inc., Markham, ON, Canada) within the corresponding restriction sites in the pME6000 plasmid (17), generating the constitutive expression vector pMCG21. All primers were purchased from Alpha DNA (Montreal, QC, Canada).

### Recombinant strains construction for BtaR2 expression

The pME6000 and pMCG21 constitutive expression vectors were introduced in *B. thailandensis* E264 strains by electroporation. Briefly, bacterial cultures were grown to an OD_600_ = 1.0, pelleted by centrifugation, and washed several times with 1 mL of sterile water. The pellets were concentrated 100-fold in 100 μL of sterile water and electroporated using a 1 mm gap disposable electroporation cuvette at 1.8 kV with an electroporator 2510 (Eppendorf Scientific, Westbury, NY, USA). Cells were outgrown for 1 hr in 1 mL Lysis Broth (LB) (Alpha Biosciences, Inc., Baltimore, MD, USA) at 37°C and plated on Tc selective media.

### Reporter strains construction

The mini-CTX-*btaI*1-*lux*, mini-CTX-*btaI*2-*lux*, and mini-CTX-*btaI*3-*lux* transcriptional reporters were integrated into the chromosome of *B. thailandensis* E264 strains through conjugation with *E. coli* χ7213 followed by selection with Tc. Successful chromosomal insertion of the *btaI*1-*lux*, *btaI*2-*lux*, and *btaI*3-*lux* plasmids was confirmed by PCR using appropriate primers.

### LC-MS/MS quantification of AHLs

The concentration of AHLs was determined from cultures of *B. thailandensis* E264 at different times during bacterial growth by liquid chromatography coupled to tandem mass spectrometry (LC-MS/MS). The samples were prepared and analyzed as described previously (18). 5,6,7,8-tetradeutero-4-hydroxy-2-heptylquinoline (HHQ-d4) was used as an internal standard. All experiments were performed in triplicate and carried out at least twice independently.

**Measurement of the *btaI*1-*lux***, ***btaI*2-*lux***, **and *btaI*3-*lux* reporters’ activity**

Expressions from the promoter regions of *btaI*1, *btaI*2, or *btaI*3 were quantified by measuring the luminescence of *B. thailandensis* E264 cultures carrying the corresponding chromosomal reporters as formerly described (13). Overnight bacterial cultures were diluted in TSB to an initial OD_600_ = 0.1 and incubated as indicated above. The luminescence was regularly determined from culture samples using a multi-mode microplate reader (Cytation^TM^ 3, BioTek Instruments, Inc., Winooski, VT, USA) and expressed in relative light units per culture optical density (RLU/OD_600_). All experiments were performed with three biological replicates and repeated at least twice.

### Quantitative reverse-transcription PCR experiments

Total RNA of *B. thailandensis* E264 cultures at an OD_600_ = 4.0 was extracted with the PureZOL RNA Isolation Reagent (Bio-Rad Laboratories, Mississauga, ON, Canada) and treated twice with the TURBO DNA-free^TM^ Kit (Ambion Life Technologies, Inc., Burlington, ON, Canada), according to the manufacturer’s instructions. Extractions were done on three different bacterial cultures. Quality and purity controls were confirmed by agarose gel electrophoresis and UV spectrophotometric analysis, respectively. cDNA synthesis was performed using the iScript^TM^ Reverse Transcription Supermix (Bio-Rad Laboratories) and amplification was accomplished on a Corbett Life Science Rotor-Gene® 6000 Thermal Cycler, using the SsoAdvanced^TM^ Universal SYBR® Green Supermix (Bio-Rad Laboratories), according to the manufacturer’s protocol. The reference gene was *ndh* (19). The *ndh* gene displayed a stable expression under the different genetic contexts tested. All primers used for cDNA amplification are presented in **Table S4**. Genes expression differences between *Burkholderia thailandensis* E264 strains were calculated using the 2^(-ΔΔ(CT))^ formula (20). A threshold of 0.5 was chosen as significant. For experiments with additions of AHLs, cultures were supplemented or not with 10 μM C_8_-HSL, 3OHC_10_-HSL, and 3OHC_8_-HSL (Sigma-Aldrich Co., Oakville, ON, Canada) from stocks prepared in HPLC-grade acetonitrile. Acetonitrile only was added in controls. All experiments were performed in triplicate and carried out at least twice independently.

### Data analysis

Unless otherwise stated, data are reported as mean +/-standard deviation (SD). Statistical analyses were performed with the R software version 3.3.3 (http://www.R-project.org.) using one-way analysis of variance (ANOVA). Probability values less than 0.05 were considered significant.

## Results

### The QS-1 and QS-2 gene clusters of *B. thailandensis* each carry an *rsaM* homologue

The *btaI*1 (*BTH_II1512*) and *btaR*1 (*BTH_II1510*) genes, respectively encoding the BtaI1 AHL synthase and the BtaR1 transcriptional regulator of the *B. thailandensis* E264 QS-1 system, are separated by the *BTH_I1511* gene that codes for an hypothetical protein conserved in the *Burkholderia* genus (7, 10, 11, 21-23). This hypothetical protein of 147 amino acids is similar to RsaM-like proteins and displays 35.8% identity with the negative AHL biosynthesis modulatory protein RsaM of the phytopathogen *P. fuscovaginae* UPB0736 (**Fig. S1A**). Interestingly, another *rsaM* homologue, encoding a hypothetical protein of uncharacterized function, is present on the genome of *B. thailandensis* E264 between the QS-2 system *btaI*2 (*BTH_II1227*) and *btaR*2 (*BTH_II1231*) genes that code for the LuxI-type synthase BtaI2 and the LuxR-type transcriptional regulator BtaR2, respectively. This hypothetical protein of 135 amino acids encoded by the *BTH_II1228* gene is 32.4% identical to RsaM (**Fig. S1A**).

Since the *BTH_II1511* and *BTH_II1228* genes are directly adjacent to *btaI*1 and *btaI*2 in the genome of *B. thailandensis* E264, respectively, and transcribed in the same direction (**Fig. S1B**), we wondered whether they could be co-transcribed. *BTH_II1228* is indeed predicted to be arranged in operon with *btaI*2 (www.burkholderia.com). According to our transcriptomic analyses (S. Le Guillouzer, M.-C. Groleau, F. Mauffrey, R. Villemur, and E. Déziel, unpublished data), neither *BTH_II1511*, nor *BTH_II1228* are co-transcribed with *btaI*1 and *btaI*2, respectively (**Fig. S1B**), as confirmed by reverse-transcription PCR (RT-PCR) experiments (data not shown).

The function of the putative proteins encoded by the *BTH_II1511* and *BTH_II1228* genes is unknown. While *BTH_II1228* is located within a cluster responsible for bactobolin biosynthesis (14, 16, 24), its involvement was actually not demonstrated. To determine whether *BTH_II1511* and *BTH_II1228* are functionally similar to the RsaM-encoding gene of *P. fuscovaginae* UPB0736, which was described as an important repressor of AHLs biosynthesis (8), we investigated the impact of these genes on the production of the predominant AHLs detected in *B. thailandensis* E264. *B. thailandensis* E264 produces 3OHC_10_-HSL and, to lesser extents, C_8_-HSL and 3OHC_8_-HSL (13-16). The total concentration of these AHLs was measured at various time intervals of the bacterial growth by LC-MS/MS in the *B. thailandensis* E264 wild-type and in the null mutants *BTH_II1511-* and *BTH_II1228-*. The mutants both overproduce AHLs when compared to the wild-type strain (**Fig. 1**). Interestingly, the impact of *BTH_II1511* on total AHL biosynthesis was more pronounced than the effect of *BTH_II1228* (**Fig. 1**). Of note, the *BTH_II1511-* mutant displays an initially delayed growth phenotype (**Fig. 1**). These observations indicate that the hypothetical proteins encoded by the *BTH_II1511* and *BTH_II1228* genes constitute, as for RsaM in *P. fuscovaginae* UPB0736, negative regulators of AHL biosynthesis and were thus designated RsaM1 and RsaM2, respectively.

**Figure 1.**
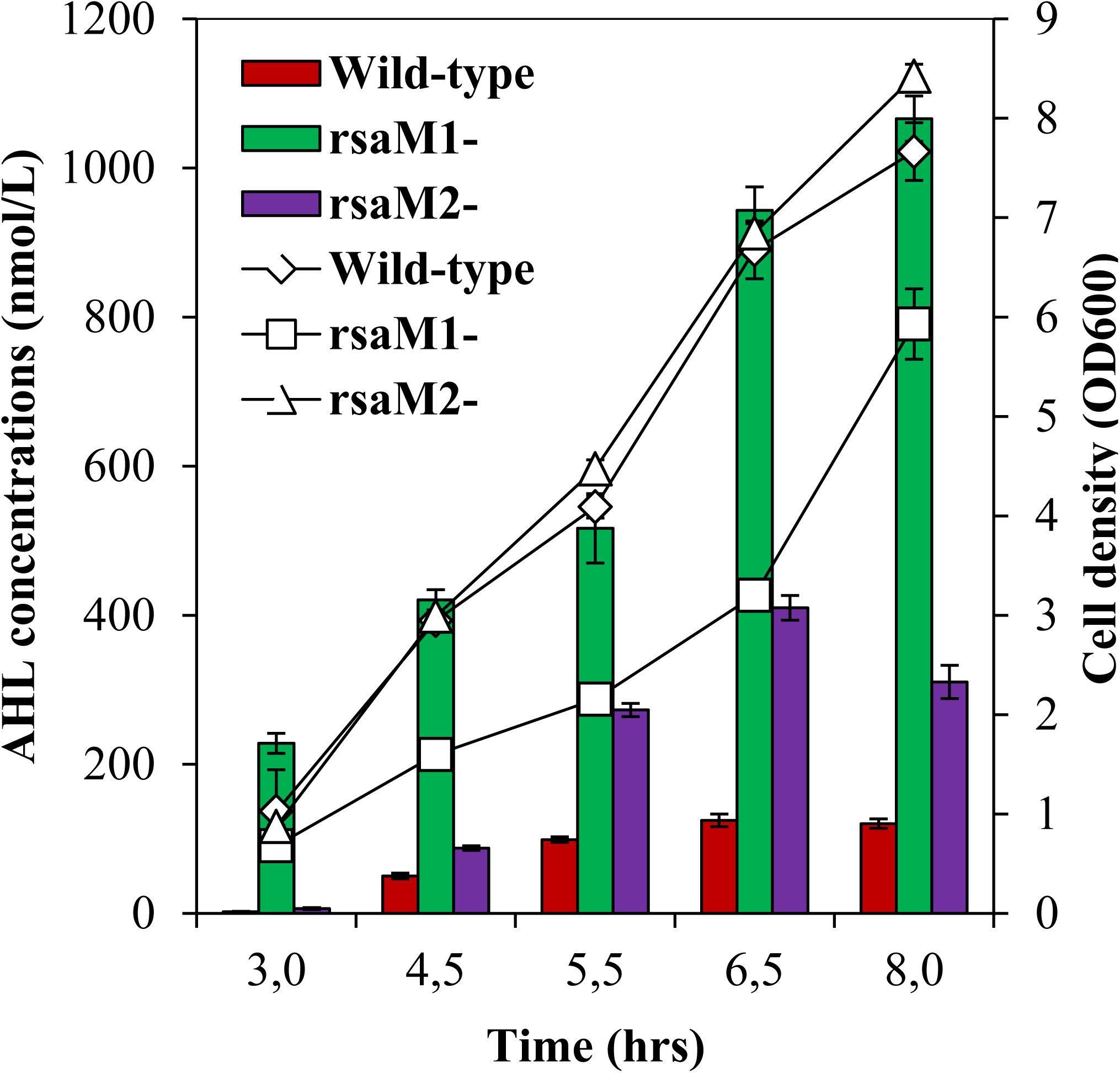
The biosynthesis of AHLs is increased in the *rsaM1* and *rsaM2* knockout mutants. Total AHLs (3OHC_10_-HSL + C_8_-HSL + 3OHC_8_-HSL) production (bars) was monitored by LC-MS/MS at various times during growth (lines) in cultures of the *B. thailandensis* E264 wild-type strain and its *rsaM*1- and *rsaM*2-mutants. The error bars represent the standard deviation of the average for three replicates.

### RsaM1 and RsaM2 are repressors of, respectively, QS-1 and QS-2 systems

To determine the effect of RsaM1 and RsaM2 on the QS-1, QS-2, and QS-3 systems, we measured the respective production of C_8_-HSL, 3OHC_10_-HSL, and 3OHC_8_-HSL in the wild-type strain E264 and in the *rsaM*1- and *rsaM*2-mutants throughout the bacterial growth phases. To gain additional insights, we also monitored expression of the AHL synthase-coding genes *btaI*1, *btaI*2, and *btaI*3 in the same backgrounds using chromosomal transcriptional fusion reporters.

We observe a dramatic overproduction of C_8_-HSL in the *rsaM*1-mutant when compared to the wild-type strain, indicating that RsaM1 negatively affects the biosynthesis of this AHL (**Fig. 2A**). Expression of the *btaI*1 gene, (14) was accordingly enhanced in the absence of RsaM1, suggesting that it is repressed by RsaM1 (**Fig. 2C**). Strikingly, the impact of RsaM1 on C_8_-HSL biosynthesis (approximately 200 fold) was more important than its effect on *btaI*1 expression (approximately 2-fold), implying that RsaM1 might also intervene at post-transcriptional levels. Additionally, C_8_-HSL concentrations also augmented in the *rsaM*2-mutant, in comparison with the wild-type strain, from the stationary phase (OD_600_ ∼8.0) (**Fig. 2B**). However, no discernible difference in *btaI*1 transcription was detected in the absence of RsaM2 (**Fig. 2C**). Altogether, these data suggest that RsaM1 represses the production of C_8_-HSL by controlling *btaI*1 transcription, including putative post-transcriptional modulations as well, whereas the negative impact of RsaM2 on C_8_-HSL biosynthesis appears to not result from *btaI*1 regulation.

**Figure 2.**
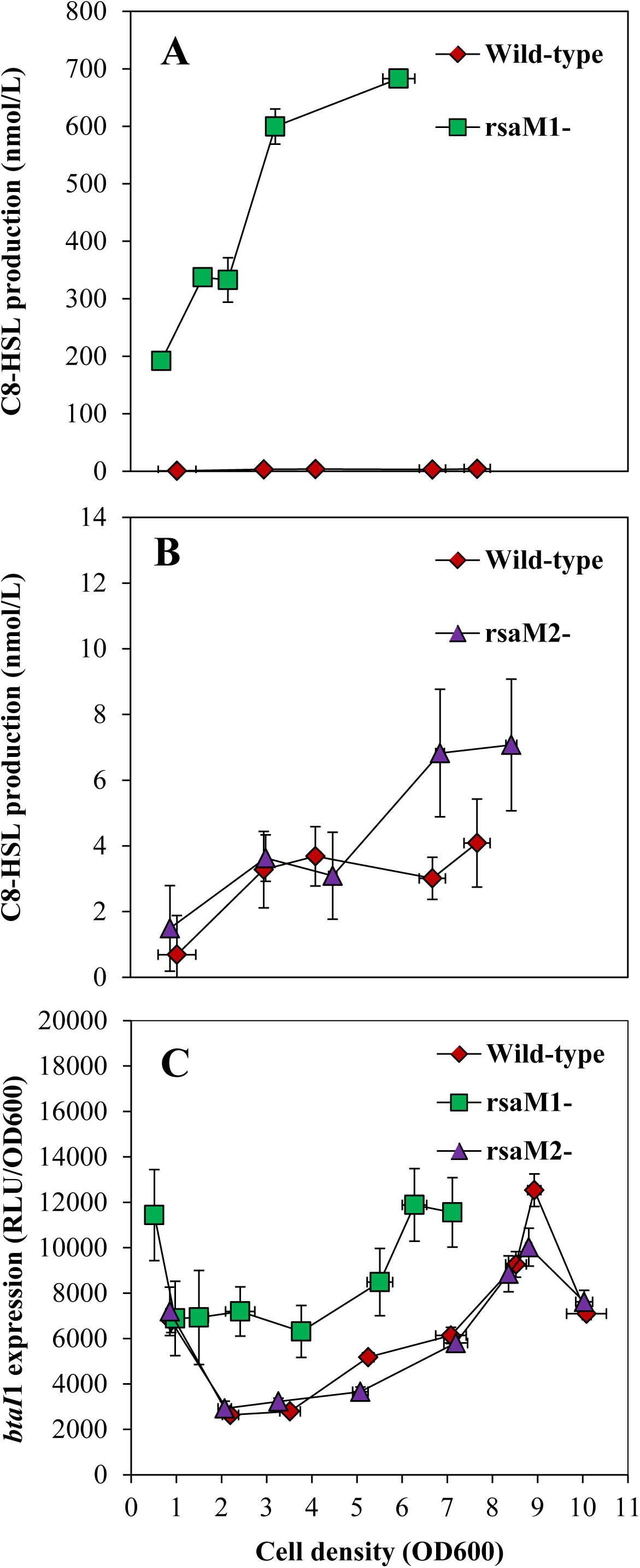
C_8_-HSL biosynthesis and expression from the *btaI***1 promoter in the wild-type and the *rsaM*1- and *rsaM*2-mutant strains of *B. thailandensis* E264**. The production of C_8_-HSL was quantified using LC-MS/MS at various times during growth in cultures of the wild-type and of (A) the *rsaM*1- and (B) *rsaM*2-mutant strains of *B. thailandensis* E264. The error bars represent the standard deviation of the average for three replicates. (C) The luminescence of the chromosomal *btaI*1-*lux* transcriptional fusion was monitored in cultures of the *B. thailandensis* E264 wild-type strain and of the *rsaM*1- and *rsaM*2-mutants. The luminescence is expressed in relative light units per culture optical density (RLU/OD_600_).

The levels of 3OHC_10_-HSL, as well as expression of the *btaI*2 gene, were unaffected in the absence of RsaM1 (**Fig. 3**). Thus, neither the production of 3OHC_10_-HSL, nor *btaI*2 transcription is under RsaM1 control. Interestingly, 3OHC_10_-HSL concentrations were strongly increased in the *rsaM*2-mutant throughout both the exponential and stationary phases, indicating that RsaM2 negatively affects 3OHC_10_-HSL biosynthesis (**Fig. 3A**). *btaI*2 transcription was similarly upregulated in the absence of RsaM2 (**Fig. 3B),** suggesting that RsaM2 represses 3OHC_10_-HSL biosynthesis by modulating the expression of *btaI*2.

**Figure 3.**
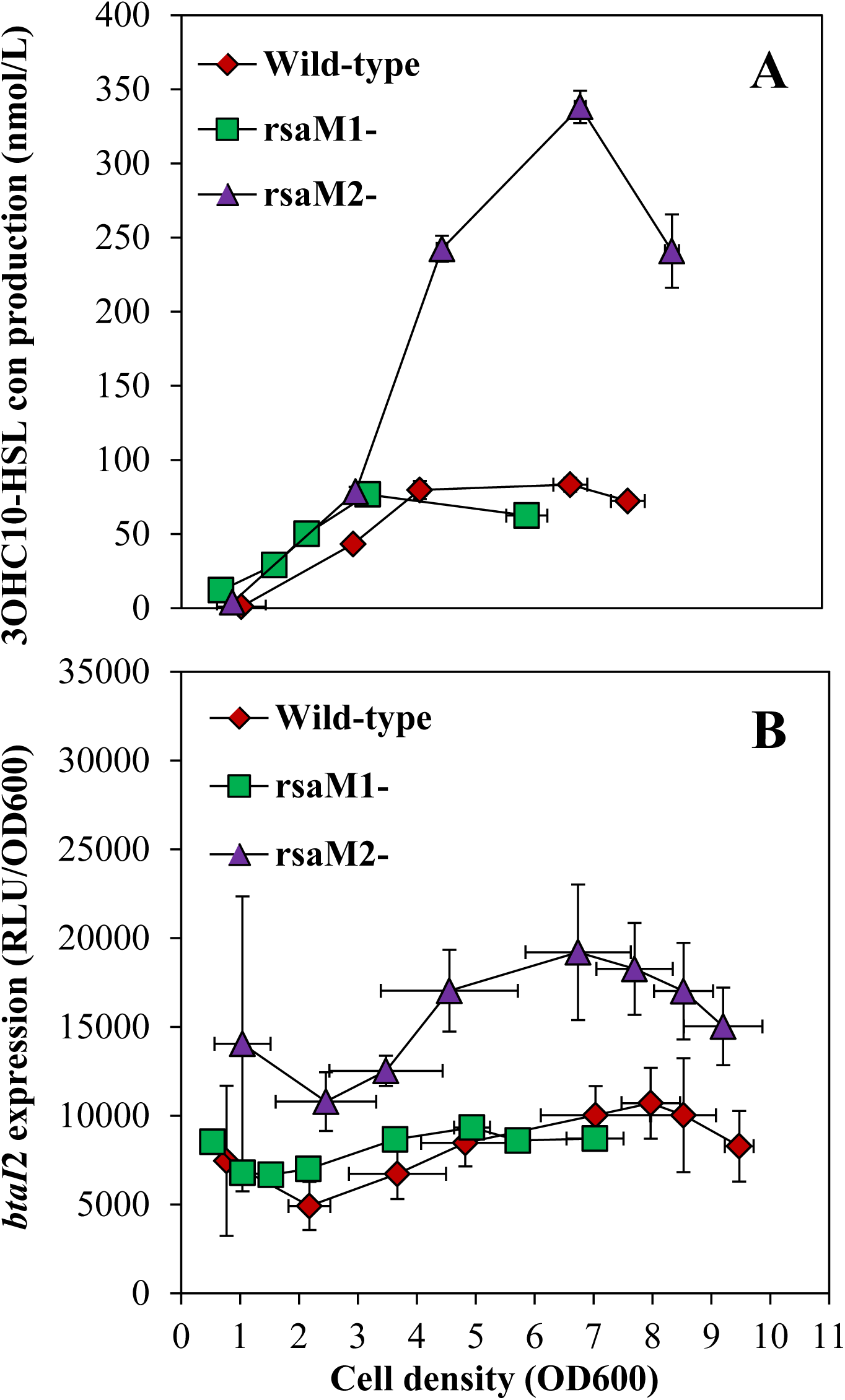
3OHC_10_-HSL biosynthesis and expression from the *btaI***2 promoter in the wild-type and the *rsaM*1- and *rsaM*2-mutant strains of *B. thailandensis* E264**. (A) The production of 3OHC_10_-HSL was quantified using LC-MS/MS at various times during growth in cultures of the wild-type and of the *rsaM*1- and *rsaM*2-mutant strains of *B. thailandensis* E264. The error bars represent the standard deviation of the average for three replicates. (B) The luminescence of the chromosomal *btaI*2-*lux* transcriptional fusion was monitored in cultures of the *B. thailandensis* E264 wild-type strain and of the *rsaM*1- and *rsaM*2-mutants. The luminescence is expressed in relative light units per culture optical density (RLU/OD_600_).

The levels of 3OHC_8_-HSL in cultures of the *rsaM*1-mutant (14) were also higher compared to the wild-type strain (**Fig. 4A**). Unexpectedly, expression of the *btaI*3 gene was not increased in the absence of RsaM1, suggesting that the negative impact of RsaM1 on 3OHC_8_-HSL production does not result from *btaI*3 regulation (**Fig. 4C**). 3OHC_8_-HSL concentrations were also increased in the *rsaM*2-mutant in comparison with the wild-type strain during the stationary phase (**Fig. 4B**), showing that the production of 3OHC_8_-HSL is repressed by RsaM2 as well. Nevertheless, no visible change in expression of *btaI*3 was noticed in the absence of RsaM2, revealing that the RsaM2-dependent regulation of 3OHC_8_-HSL biosynthesis might also not be linked to *btaI*3 (**Fig. 4C**).

**Figure 4.**
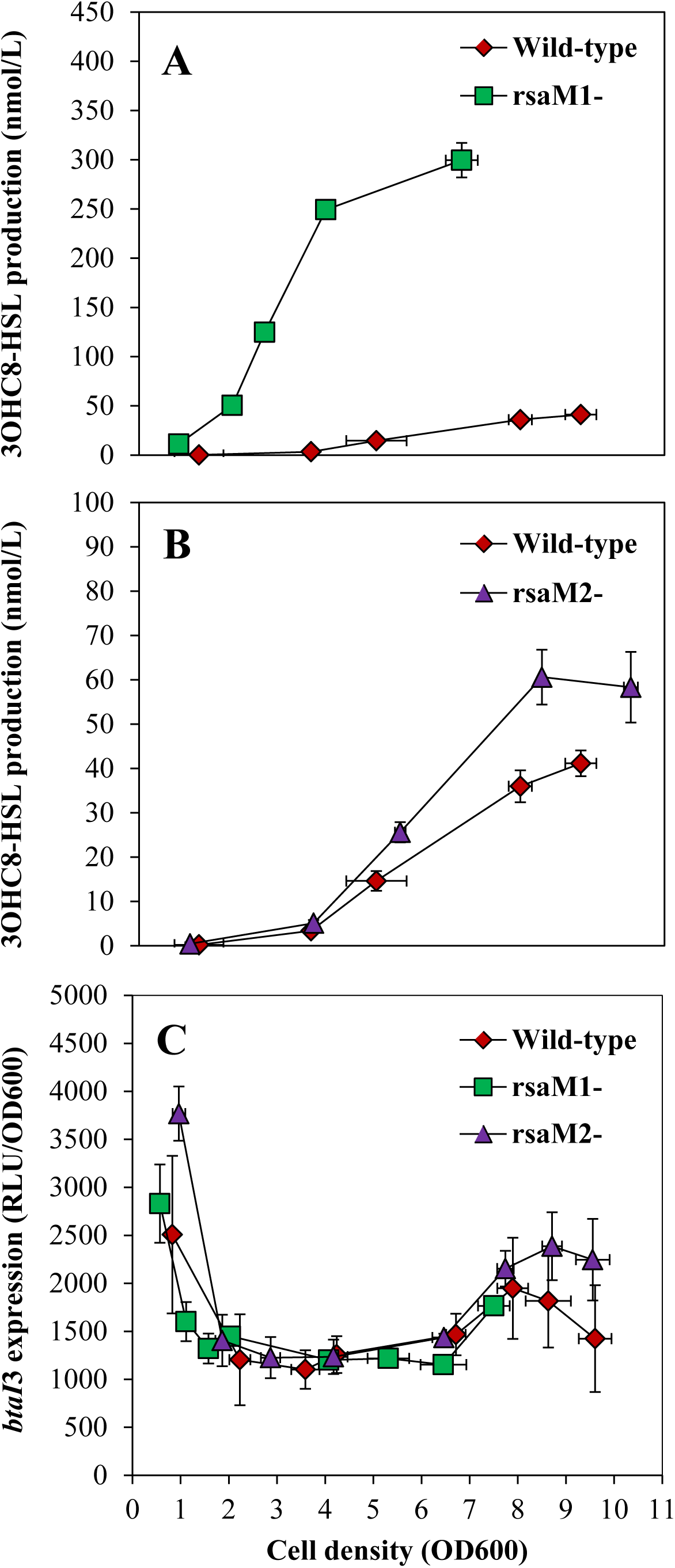
3OHC_8_-HSL biosynthesis and expression from the *btaI***3 promoter in the wild-type and the *rsaM*1- and *rsaM*2-mutant strains of *B. thailandensis* E264**. The production of 3OHC_8_-HSL was quantified using LC-MS/MS at various times during growth in cultures of the wild-type and of (A) the *rsaM*1- and (B) *rsaM*2-mutant strains of *B. thailandensis* E264. The error bars represent the standard deviation of the average for three replicates. (C) The luminescence of the chromosomal *btaI*3-*lux* transcriptional fusion was monitored in cultures of the *B. thailandensis* E264 wild-type strain and of the *rsaM*1- and *rsaM*2-mutants. The luminescence is expressed in relative light units per culture optical density (RLU/OD_600_).

While the concentrations of both C_8_-HSL and 3OHC_8_-HSL were enhanced in the *rsaM*1-mutant, the impact on the former was more important (**Figs. S2A and S2B**). Additionally, C_8_-HSL, 3OHC_10_-HSL, and 3OHC_8_-HSL levels were all increased in the absence of RsaM2, however, 3OHC10-HSL levels were more affected (**Figs. S2A and SC**). Collectively, these findings indicate that the QS-1 system is mainly repressed by RsaM1, whereas RsaM2 principally represses the QS-2 system.

### RsaM1 negatively regulates transcription of *btaR*1 but expression of the *btaR*2 gene is not modulated by RsaM2

In order to determine whether the impact of RsaM1 and RsaM2 on AHL biosynthesis also involves the BtaR1, BtaR2, and BtaR3 transcriptional regulators, (13, 15, 16), we monitored expressions of their respective encoding genes by quantitative reverse-transcription PCR (qRT-PCR) in the wild-type strain and in the *rsaM*1- and *rsaM*2-mutants during the exponential phase. Interestingly, we observed an increase in *btaR*1 transcription in the absence of RsaM1 (**Fig. 5A**), which correlates with the expression profile of *btaI*1 in this background (**Fig. 5B**). However, no variation was observed in the *rsaM*2-mutant when compared to the wild-type strain (**Fig. 5**). Thus, while expression of both *btaR*1and *btaI*1 are negatively regulated by RsaM1, RsaM2 does not impact any of the QS-1 system regulatory genes. Taken together, these results suggest that the negative impact of RsaM1 on the QS-1 system also implies regulation of *btaR*1, whereas RsaM2 might exclusively act at post-transcriptional levels. Furthermore, no discernible difference was detected in the *btaR*2 gene transcription in the *rsaM*1-mutant strain in comparison with the wild-type strain, and its expression was also unchanged in the absence of RsaM2, showing that RsaM1 nor RsaM2 modulate expression of *btaR*2 (data not shown). Consequently, while RsaM1 seems to have no effect on the QS-2 system, RsaM2-dependent regulation of the QS-2 system is apparently not linked to *btaR*2 control but might rather go through regulation of *btaI*2 transcription. Moreover, neither RsaM1 nor RsaM2 modulate the transcription of *btaR*3 (data not shown). Collectively, these observations indicate that both RsaM1 and RsaM2 do not regulate 3OHC_8_-HSL biosynthesis through the QS-3 system genes *btaR*3and *btaI*3, which encodes the BtaI3 synthase mainly responsible for 3OHC_8_-HSL production.

**Figure 5.**
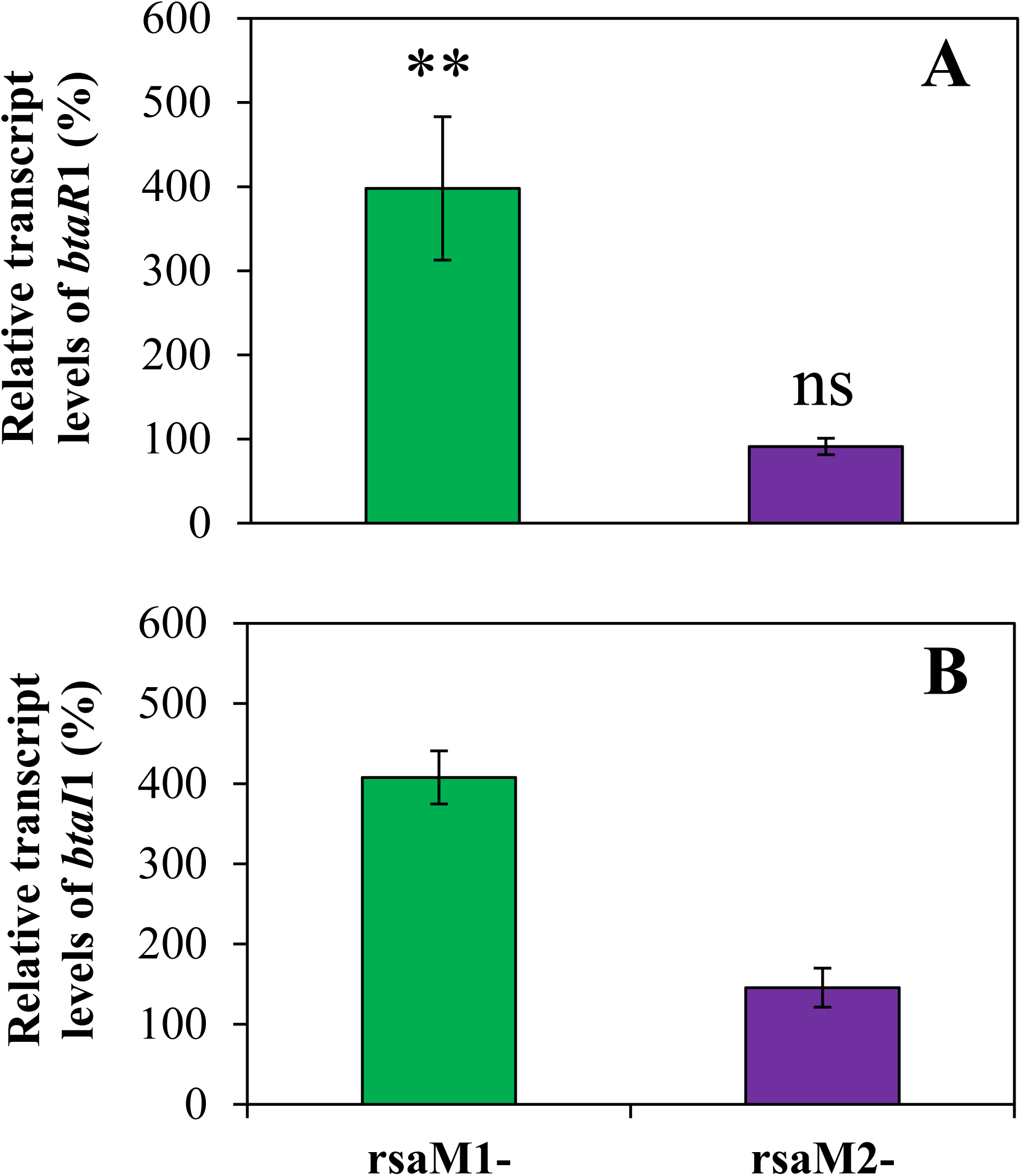
Expressions of *btaR***1 and *btaI*1 are both repressed by RsaM1**. The relative transcript levels of (A) *btaR*1 and (B) *btaI*1 were assessed by qRT-PCR experiments in cultures of the wild-type and of the *rsaM*1- and *rsaM*2-mutant strains of *B. thailandensis* E264. The results are presented as relative quantification of genes transcription compared to the wild-type normalized to 100%. The error bars represent the standard deviation of the average for three replicates. **, p < 0.01; ns, non-significant.

### The *rsaM*1 and *rsaM*2 genes are QS-controlled

Our transcriptomic sequencing analyses indicate that QS positively regulates the expression of *rsaM1* and *rsaM2* (S. Le Guillouzer, M. C. Groleau, F. Mauffrey, R. Villemur, and E. Déziel, unpublished data). Majerczyk *et al*. (15) also reported that the *rsaM*2 gene is QS-controlled, but not *rsaM*1.

In order to ascertain that *rsaM*1 is under QS control, we monitored *rsaM*1 expression by qRT-PCR in the *B. thailandensis* E264 wild-type strain and in the AHL-null Δ*btaI*1Δ*btaI*2Δ*btaI*3 mutant supplemented or not with exogenous AHLs during the exponential phase. We observed that expression of *rsaM*1 is reduced in the absence of AHLs (**Fig. 6A**), confirming that QS positively modulates *rsaM*1 transcription. Furthermore, expression of *rsaM*1 was restored to wild-type levels in the culture of the AHL-null mutant strain supplemented with either C_8_-HSL or 3OHC_8_-HSL (**Fig. 6A**). Moreover, adding 3OHC_10_-HSL to the culture of the Δ*btaI*1Δ*btaI*2Δ*btaI*3 mutant background did not significantly enhance *rsaM*1 transcription (**Fig. 6A**). To gain insights into the QS-dependent regulation of *rsaM*1, we also measured expression of *rsaM*1 in the Δ*btaR*1, Δ*btaR*2, and Δ*btaR*3 mutants *vs*. the *B. thailandensis* E264 wild-type strain during the exponential phase. While no obvious change in *rsaM*1 transcription was visible in the absence of neither BtaR2, nor BtaR3 under our conditions, expression of *rsaM*1 was decreased in the Δ*btaR*1 mutant when compared to the wild-type strain (**Fig. 6B**). Taken together, these data indicate that expression of *rsaM*1 is positively regulated by the QS-1 system and might be activated by BtaR1 in association with C_8_-HSL or 3OHC_8_-HSL, whereas BtaR2 and BtaR3 are not directly involved in the modulation of *rsaM*2 transcription.

**Figure 6.**
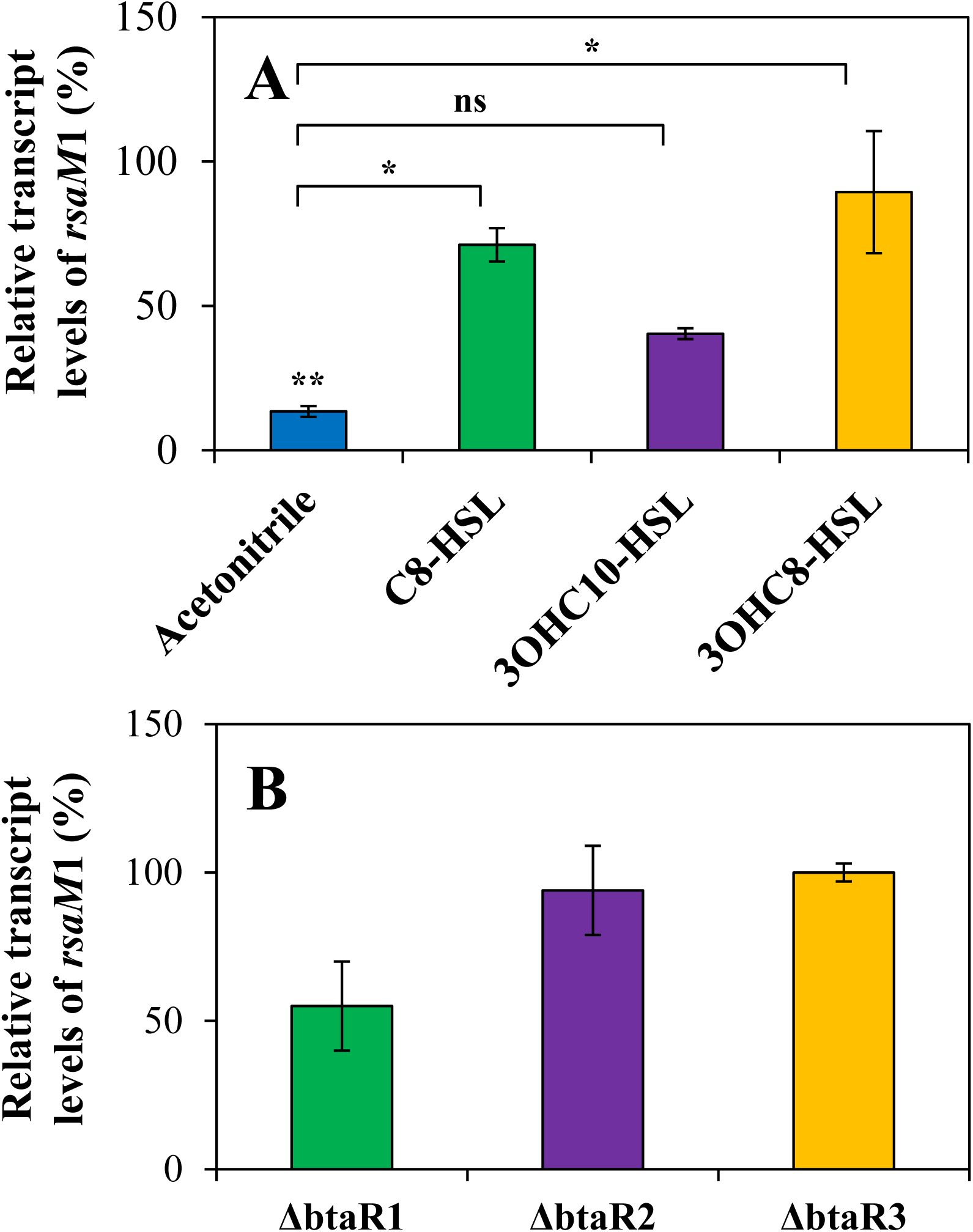
QS positively regulates *rsaM* 1 transcription. (A) The relative transcript levels of *rsaM*1 from the *B. thailandensis* E264 wild-type and its ΔbtaI1ΔbtaI2ΔbtaI3 mutant strain were estimated by qRT-PCR experiments. Cultures were supplemented with 10 μM C_8_-HSL, 3OHC_10_-HSL, or 3OHC_8_-HSL. Acetonitrile only was added in controls. The results are presented as relative quantification of genes transcription compared to the wild-type normalized to 100%. The error bars represent the standard deviation of the average for three replicates. (B) The relative transcript levels of *rsaM*1 were assessed by qRT-PCR experiments in cultures of the wild-type and of the ΔbtaR1, ΔbtaR2, and Δ*btaR*3 mutant strains of *B. thailandensis* E264. **, p < 0.01; *, p < 0.05; ns, non-significant.

Expression of *rsaM*2 was lowered in the absence of AHLs, confirming that the *rsaM*2 gene is activated by QS (**Fig. 7A**). Furthermore, *rsaM*2 transcription was restored to wild-type levels in cultures of the Δ*btaI*1Δ*btaI*2Δ*btaI*3 mutant strain supplemented with either 3OHC_10_-HSL or 3OHC_8_-HSL (**Fig. 7A**). Moreover, adding C_8_-HSL did not significantly increase *rsaM*2 transcription, revealing that the *rsaM*2 gene is not activated by C_8_-HSL (**Fig. 7A**). Interestingly, we observed that expression of *rsaM*2 was also downregulated in the Δ*btaR*2 mutant in comparison with the wild-type strain, meaning that the *rsaM*2 gene is positively controlled by BtaR2, whereas no discernible difference in *rsaM*2 transcription was detected in the absence of neither BtaR1, nor BtaR3 under the conditions of our experiments (**Fig. 7B**). Altogether, these results indicate that expression of *rsaM*2 is positively regulated by the QS-2 system and might be activated by BtaR2 in response to 3OHC_10_-HSL or 3OHC_8_-HSL, whereas BtaR1 and BtaR3 do not intervene in *rsaM*2 regulation.

**Figure 7.**
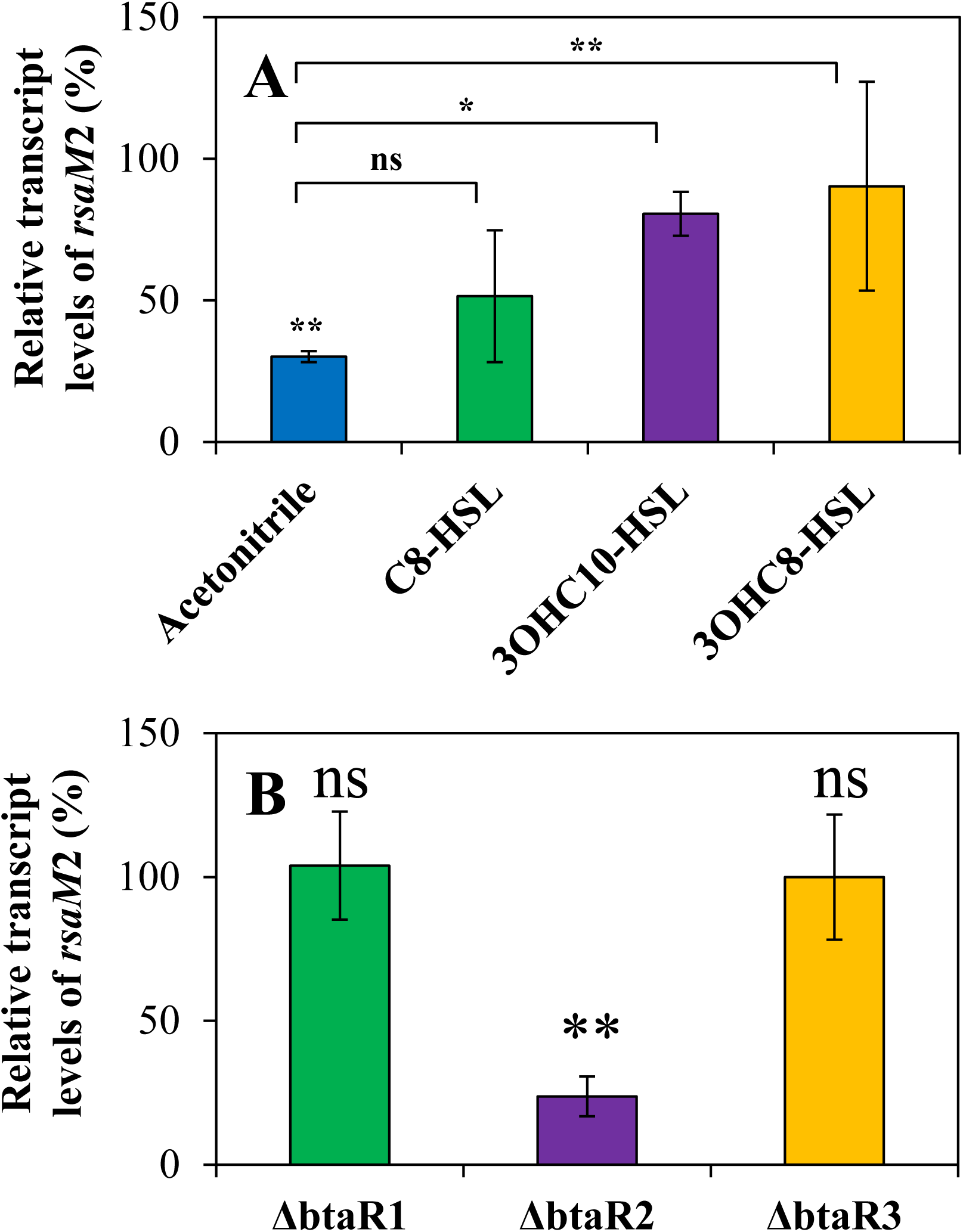
Expression of *rsaM***2 is activated by QS**. (A) The relative transcript levels of *rsaM*2 from the *B. thailandensis* E264 wild-type and its ΔbtaI1ΔbtaI2ΔbtaI3 mutant strain were monitored by qRT-PCR experiments. Cultures were supplemented with 10 μM C_8_-HSL, 3OHC_10_-HSL, or 3OHC_8_-HSL. Acetonitrile only was added in controls. The results are presented as relative quantification of genes transcription compared to the wild-type normalized to 100%. The error bars represent the standard deviation of the average for three replicates. (B) The relative transcript levels of *rsaM*2 were quantified by qRT-PCR experiments in cultures of the wild-type and of the ΔbtaR1, ΔbtaR2, and ΔbtaR3 mutant strains of *B. thailandensis* E264. **, p < 0.01; *, p < 0.05; ns, non-significant.

Collectively, these observations highlight that expression of *rsaM*1is activated by the QS-1 system, which is negatively controlled by RsaM1, whereas *rsaM*2 transcription is positively regulated by the QS-2 system, which is repressed by RsaM2, showing that these RsaM-like proteins are integrated into the QS modulatory web of *B. thailandensis* E264.

### Expression of *rsaM*1 and *rsaM*2 are negatively auto-regulated

To further explore the RsaM1 and RsaM2 molecular mechanisms of action, *rsaM*1 and *rsaM*2 expressions were assessed by qRT-PCR in the wild-type strain of *B. thailandensis* E264 and in the *rsaM*1- and *rsaM*2-mutants during the exponential phase. Expression of *rsaM*1 was increased in the *rsaM*1-mutant when compared to the wild-type strain, and the same was observed for *rsaM*2 expression in the *rsaM2-* mutant (**Figs. S3A and S3B**). However, the absence of RsaM2 had no impact on *rsaM*1 transcription (**Fig. S3A**) and *rsaM*2 transcription was unchanged in the *rsaM*1-mutant in comparison with the wild-type strain (**Fig. S3B**). Altogether, these results indicate that RsaM1 and RsaM2 repress their own expression, but are not transcriptionally hierarchically organized.

## Discussion

*B. thailandensis* E264 can synthesize C_8_-HSL, 3OHC_10_-HSL, and 3OHC_8_-HSL (13-16), with 3OHC_10_-HSL being the most abundant AHL detected during the different stages of growth. This reveals that the production of C_8_-HSL and 3OHC_8_-HSL might be under stringent control. These signaling molecules mediate the activity of three BtaR/BtaI QS systems. Here, we initiated the study of two uncharacterized genes present in the QS-1 and QS-2 gene clusters. RsaM1 and RsaM2 were shown to dramatically restrict the production of AHLs, highlighting their deep involvement in the complex organization of the multiple AHL-based QS circuits of *B. thailandensis* E264, as summarized in **Fig. 8**.

**Figure 8.**
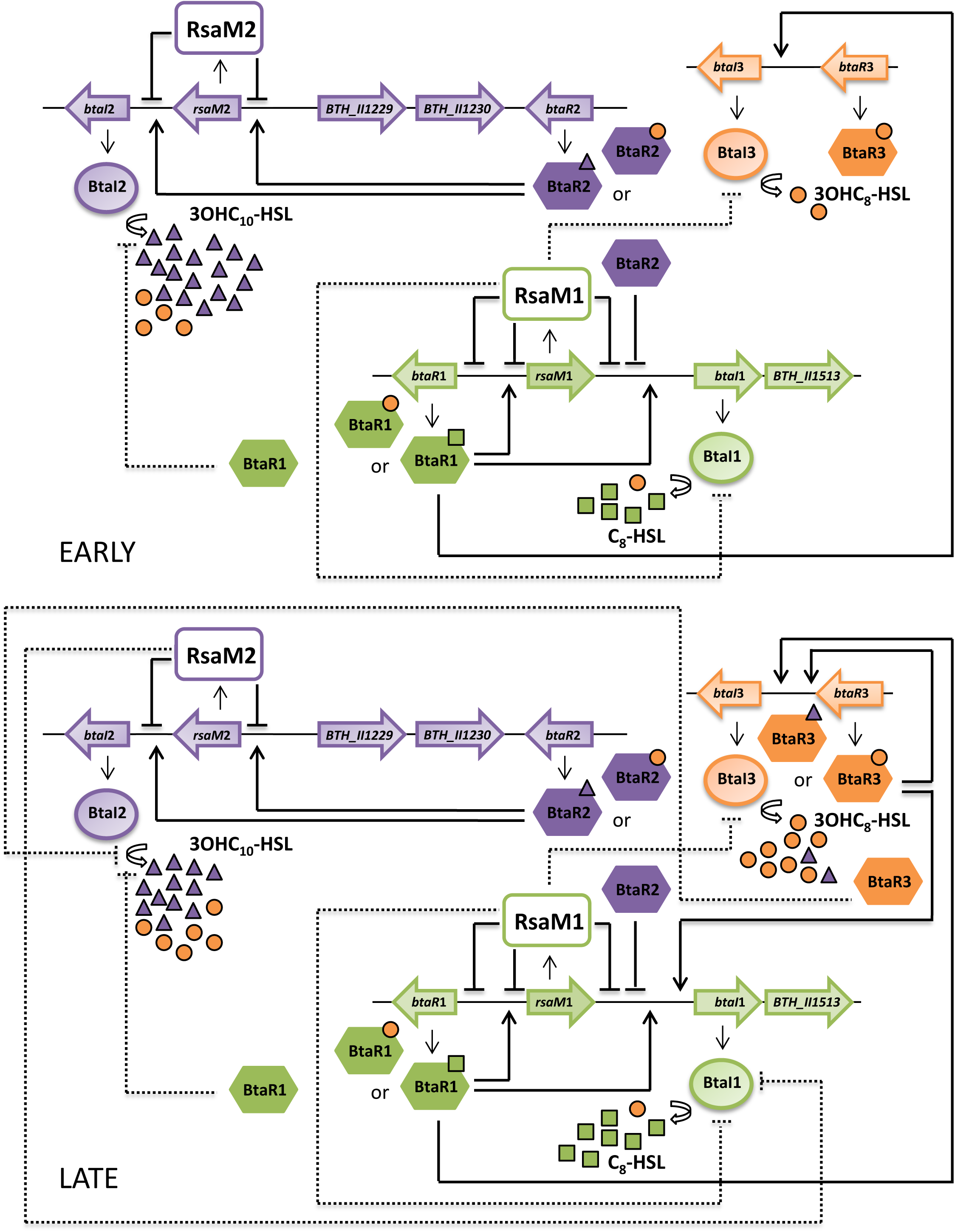
Proposed model of the QS regulatory network of *B. thailandensis* E264.

A gene conserved in the *Burkholderia* genus within the QS-1 system (7, 10, 11, 21-23) is divergently transcribed from *btaR*1 and oriented in the same direction as *btaI*1 **(Fig. 8 and Fig. S1B**). This gene encodes an hypothetical protein, homologous to the negative AHL biosynthesis modulatory protein RsaM originally identified in the plant pathogen *P. fuscovaginae* UBP0736 (8, 9), hence designated RsaM1. In *B. cenocepacia* H111, the homologue *Bc*RsaM was described as an important repressor of the CepI/CepR QS system, and proposed to inhibit the production of C_8_-HSL by regulating the activity and/or stability of the LuxI-type synthase CepI and the LuxR-type transcriptional regulator CepR, as well as the orphan LuxR-type transcriptional regulator CepR2 (10, 11). While the transcription of the QS *cepI*, *cepR*, and *cepR*2 genes were shown to be lowered in the *rsaM*-mutant of *B. cenocepacia* H111 (10), *btaI*1 and *btaR*1 expressions were both increased in our RsaM1 of *B. thailandensis* E264, correlating with the accumulation of C_8_-HSL observed in this background. Consequently, RsaM1 could repress the transcription of *btaI*1 and *btaR*1, suggesting that its mode of action in *B. thailandensis* differs from that of *Bc*RsaM. Still, the impact of RsaM1 on C_8_-HSL biosynthesis was dramatically higher than its effect on *btaI*1 expression (**Figs. 2A and 2C**), which hints that RsaM1 might also act at post-transcriptional and/or post-translational levels, as proposed for *Bc*RsaM. Thus, RsaM1 could repress as well the expression of *btaI*1 and *btaR*1 directly or indirectly, for instance by modulating the synthase activity and/or stability of BtaI1 (14), or by controlling the functionality of BtaR1. Clearly, while BtaR1 is considered the principal regulator of *btaI*1 expression, RsaM1 plays a major role in modulating the production of C_8_-HSL.

We recently determined that expression of *btaI*1 is positively controlled by BtaR1, as well as activated by the presence of C_8_-HSL. The accumulation of this AHL detected in the Δ*btaR*1 mutant background is not directly induced by the impact of BtaR1 on *btaI*1 expression, but could involve additional transcriptional and/or post-transcriptional regulators under its control (13). Interestingly, we observed that *rsaM*1 expression is positively controlled by BtaR1/C_8_-HSL (**Fig. 6**), suggesting that the overproduction of C_8_-HSL detected in the Δ*btaR*1 mutant strain is indirectly induced by BtaR1 through RsaM1. This also reveals that the QS-1 system is negatively auto-regulated, maybe counteracting the positive feedback loop mediated by BtaR1/C_8_-HSL for C_8_-HSL biosynthesis, and thus be necessary to maintain the production of this AHL within optimal and adequate levels according to specific environmental conditions, as previously suggested for the RsaL repressor in *P. aeruginosa* PAO1 (23, 25). In agreement with the finding that *rsaM* is positively and directly regulated by CepR in *B. cenocepacia* J2315 (10, 26), we found a putative *lux*-box sequence in the promoter region of the *rsaM*1 gene (**Fig. S1C**), suggesting that *rsaM*1 could also be directly under BtaR1 control in *B. thailandensis* E264. Nevertheless, *rsaM*1 transcription was not completely abolished in the absence of BtaR1, indicating that the QS-dependent regulation of the *rsaM*1 gene is be more complex and requires further investigation.

Strikingly, the absence of RsaM1 was associated with a growth delay of the mutant (**Fig. 1**), correlating with aggregation of cells. This could be linked to the high levels of C_8_-HSL produced in this background and thus over-activation of phenotypes controlled by the QS-1 system such as auto-aggregation and oxalate production (14, 15, 27, 28). Thus, RsaM1 could be necessary for the regulation of phenotypic traits under BtaR1/C_8_-HSL control, requiring substantial concentrations of C_8_-HSL under specific environmental conditions, as suggested for the RsaL repressor in *B. kururiensis* M130 (29).

Interestingly, RsaM1 also represses the production of 3OHC_8_-HSL (**Fig. 4A),** an AHL synthesized by both BtaI2 and BtaI3 (14, 16). Because no overproduction of 3OHC_10_-HSL, the main AHL produced by BtaI2 (16), was observed in the *rsaM*1-mutant background (**Fig. 3A**), we assume that the QS-2 system is not influenced by RsaM1. Thus, while RsaM1 seems to act on 3OHC_8_-HSL by modulating the QS-3 system, although no impact on expression of *btaI*3 and *btaR*3 was observed, we propose that its negative impact is going through an effect on the activity and/or stability of the BtaI3 synthase. This would add another regulatory layer linking the QS-1 and QS-3 systems in *B. thailandensis* E264, already shown to be hierarchically organized at the transcriptional level (13, 15). Additionally, it is conceivable that concentrations of 3OHC_8_-HSL produced by BtaI1 in the wild-type strain are below our detection limit and that these levels become detectable in the QS-1-boosted *rsaM*1-mutant background. Indeed, we previously reported production of both C_8_-HSL and traces amounts of 3OHC_8_-HSL by the same synthase in a *Burkholderia* strain from the *cepacia* complex (30), and the *B. pseudomallei* KHW BpsI and *B. mallei* ATCC 23344 BmaI1 synthases were both shown to produce low levels of 3OHC_8_-HSL in addition to C_8_-HSL, (31, 32). Additionally, while we demonstrated that *rsaM*1 expression is controlled by BtaR1 in response to C_8_-HSL, activation of this gene by 3OHC_8_-HSL might also involve BtaR1 (**Fig. 6**). This is further supported by the fact that the *B. pseudomallei* KHW BpsR and *B. mallei* ATCC 23344 BmaR1 transcriptional regulators were reported to specifically respond to both C_8_-HSL and 3OHC_8_-HSL, with C_8_-HSL eliciting the strongest response (31, 32). This is in agreement with the finding that the BtaR1-controlled genes identified in transcriptomic analyses were generally affected by both C_8_-HSL and 3OHC_8_-HSL (15). This would also explain why 3OHC_8_-HSL levels increased in the absence of BtaR1, following the same kinetic as C_8_-HSL (13), indicating that RsaM1 could play a part in the accumulation of this AHL detected in the Δ*btaR*1 mutant background as proposed for C_8_-HSL. Nevertheless, additional experiments will be necessary to confirm the production of 3OHC_8_-HSL by BtaI1.

Importantly, the fact that the production of both C_8_-HSL and 3OHC_8_-HSL, but not 3OHC10-HSL, is stringently repressed by RsaM1, is consistent with the recent finding that C_8_-HSL and 3OHC_8_-HSL are synthesized at low levels throughout the different stages of growth (13), and could reasonably justify the predominance of 3OHC_10_-HSL in *B. thailandensis* E264.

Interestingly, an *rsaM* homologue encoding a hypothetical protein designated RsaM2 is also found directly adjacent to *btaI*2 and transcribed in the same direction **(Fig. 8 and Fig. S1B**). We recently corroborated that *btaI*2 transcription is principally activated by BtaR2 in association with 3OHC_10_-HSL (13). While BtaI2 is mainly responsible for the production of this AHL (16), and we observed that 3OHC_10_-HSL biosynthesis is almost completely abolished in the Δ*btaI*2 mutant (**Fig. S4B**), we demonstrated that the production of 3OHC_10_-HSL is positively regulated by BtaR2 (13). Furthermore, our results show that RsaM2 represses the biosynthesis of 3OHC_10_-HSL, as well as the expression of *btaI*2, indicating that RsaM2-dependent regulation of the QS-2 system is apparently not linked to *btaR*2 control but might rather go through regulation of *btaI*2 transcription. Since 3OHC_10_-HSL biosynthesis and expression of *btaI*2 were similarly affected by RsaM2 (**Fig. 3**), the activity and/or stability of BtaI2 might not be altered by RsaM2. However, we do not exclude that RsaM2 represses 3OHC_10_-HSL biosynthesis by controlling the functionality of BtaR2. Moreover, we observed that expression of *rsaM*2 is activated by the QS-2 system (**Fig. 7**). Consequently, while expression of *btaI*2 is directly activated by BtaR2, we demonstrated that BtaR2 also represses *btaI*2 transcription indirectly through RsaM2 control. This homeostatic control of the QS-2 system appears similar to what proposd for the RsaL repressor in *P. aeruginosa* PAO1 (23, 25).

We also highlighted that the production of 3OHC_8_-HSL is negatively regulated by RsaM2 (**Fig. 4B**), and because we determined that expressions of neither *btaI*3, nor *btaR*3 are under RsaM2 control, we must conclude that RsaM2 does not influence this AHL by modulating the QS-3 system. Interestingly, the production of 3OHC_8_-HSL is repressed by RsaM1 as well (**Fig. 4A**). This could also explain why inactivating either *rsaM*1 or *rsaM*2 does not result in *btaI*3 activation; investigating the expression of this gene in a double *rsaM*1-*rsaM*2-mutant would be necessary to determine the precise regulatory mechanism directing 3OHC_8_-HSL biosynthesis through these RsaM-like proteins. Nevertheless, since BtaI2 produces this AHL in addition to 3OHC_10_-HSL, albeit to a lesser extent (16), we suppose that the negative impact of RsaM2 on the production of 3OHC_8_-HSL, as the RsaM2-dependent regulation of 3OHC_10_-HSL biosynthesis, is linked to *btaI*2 regulation. Remarkably, we noticed that the production of 3OHC_10_-HSL is repressed by RsaM2 from the exponential phase, whereas 3OHC_8_-HSL biosynthesis is repressed by RsaM2 from the stationary phase. This is consistent with our hypothesis that 3OHC_8_-HSL is produced by BtaI2 at the expense of 3OHC_10_-HSL (13), meaning that BtaI2 mainly synthesizes 3OHC_10_-HSL during the logarithmic growth, whereas it principally synthesizes 3OHC_8_-HSL during the stationary phase, further confirmed by 3OHC_8_-HSL being produced in the stationary phase, but almost not in the logarithmic growth when BtaI3 expression is very low (**Fig. S4C**).

We confirmed that the production of C_8_-HSL is completely abolished in the absence of BtaI1, indicating that this AHL is exclusively produced by this synthase (**Fig. S4A**); indeed, no other synthase than the homologs BpsI and BmaI1 have been associated with the production of this AHL in *B. pseudomallei* KHW and *B. mallei* ATCC 23344, respectively (31, 32). Consequently, it is not clear how C_8_-HSL biosynthesis is repressed by RsaM2 when no matching overexpression of *btaI*1 is observed in the absence of RsaM2 (**Figs. 2B and 2C**). The QS-1 and QS-2 systems were recently found to be sequentially organized (13), and we indeed confirmed that BtaR2 negatively modulates the production of C_8_-HSL by repressing the expression of *btaI*1 (**Figs. S5 and S6**). We propose that the negative impact of RsaM2 on the production of C_8_-HSL might involve post-transcriptional regulation, underscoring an additional regulatory layer connecting the QS-1 and QS-2 systems in *B. thailandensis* E264

We also demonstrated that RsaM1 and RsaM2 repress their own transcription (**Fig. S3**). Negative auto-regulation of these RsaM-like proteins could be necessary to maintain AHLs at appropriate levels depending on particular environmental conditions, and might further contribute to the correct timing of the QS-1, QS-2, and QS-3 systems response. Since RsaM1 and RsaM2 had no impact on *rsaM*2 and *rsaM*1 expression, respectively, we hypothesize that these RsaM homologues act on the biosynthesis of common AHLs independently of each other, thus ensuring the specificity of the regulation of the QS-1, QS-2, and QS-3 systems.

**Figure 8** shows the proposed interactions between the QS-1, QS-2, and QS-3 systems and the RsaM homologues RsaM1 and RsaM2. We previously showed that the first system activated in *B. thailandensis* E264 is QS-2 with 3OHC_10_-HSL production starting earlier and at higher levels than the other AHLs (13). BtaR2 activates *rsaM*2 expression. RsaM2 has a negative impact on 3OHC_10_-HSL. We also proposed that the QS-2 system has a negative impact on the QS-1 system (13) and we believe this effect goes through RsaM1. However, there is no effect on *rsaM*1 expression in a Δ*btaR*2 mutant, which adds to our previous hypothesis that BtaR2 acts on the transcription of *btaI*1. Our qRT-PCR experiments indicate that BtaR1 activates expression of *rsaM*1, which controls negatively the biosynthesis of C_8_-HSL. Since our previous results show that a Δ*btaR*1 mutant produces higher levels of C_8_-HSL than the wild-type with no matching overexpression of *btaI*1, it is possible in the inactivation of *btaR*1 affects *rsaM*1 expression and thus BtaI1 activity. The effect of the QS-1 system on the QS-3 system was also formerly detailed (13).

## Conclusion

We reported that the QS-1, QS-2, and QS-3 systems are hierarchically and homeostatically arranged in *B. thailandensis* E264 and we also observed that these QS systems are integrated into an intricate network, including additional unspecified transcriptional and/or post-transcriptional regulators (13). The present study uncovers the crucial role of the two newly identified RsaM homologues designated RsaM1 and RsaM2 in the modulation of AHL signaling (**Fig. 8**). We demonstrated that the QS-1 system is mainly repressed by RsaM1, whereas RsaM2 principally represses the QS-2 system. Additionally, these AHL biosynthesis regulatory proteins were shown to be an integral part of the QS modulatory circuitry, contributing to the temporal expression of the multiple AHL-based QS circuits of *B. thailandensis* E264. The precise underlying molecular mechanism of action of RsaM-like proteins remains currently unknown and has to be further investigated in the future given their importance in the regulation of QS-controlled genes in the *Burkholderia* genus and other Proteobacteria (7-11, 21-23).

**Funding information**

This study was supported by Canadian Institutes of Health Research (CIHR) Operating Grants MOP-97888 and MOP-142466 to Eric Déziel. Eric Déziel holds the Canada Research Chair in Sociomicrobiology. The funders had no role in study design, data collection and interpretation, or the decision to submit the work for publication.

## Acknowledgments

Thank you to Everett Peter Greenberg (Department of Microbiology, University of Washington School of Medecine, Seattle, WA, USA) for providing *B. thailandensis* E264 strains. Special thanks to Sylvain Milot and François D’Heygere for their technical help.

## Legends for supplemental material

**Figure S1.**
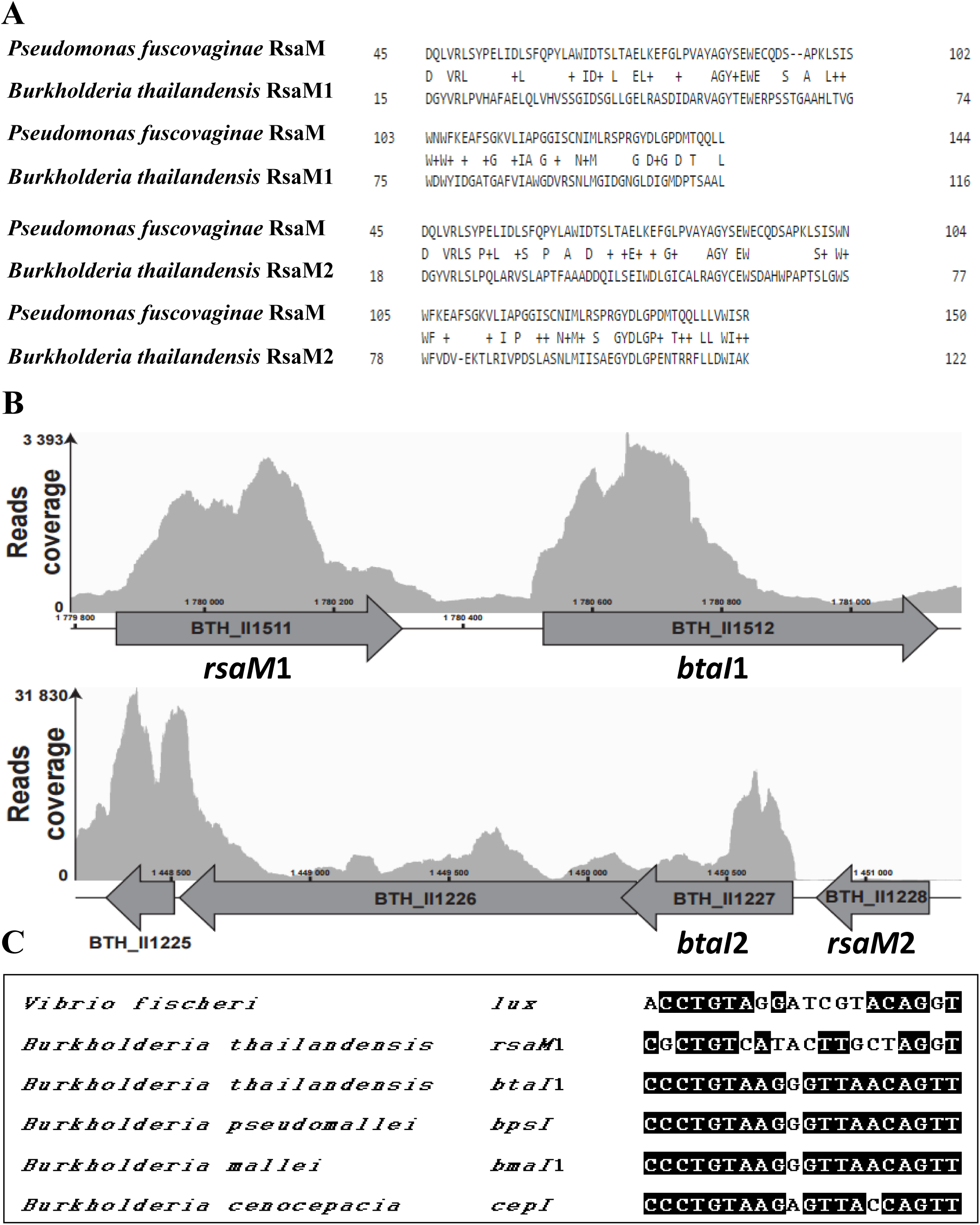
*B. thailandensis* possesses two conserved RsaM-like proteins designated RsaM1 and RsaM2. (A) Sequence alignment of the RsaM1 and RsaM2 proteins of *B. thailandensis* E264 with the *P. fuscovaginae* UPB0736 RsaM homologue. (B) Genetic arrangement of *rsaM1* and *rsaM2* with *btaI*1 and *btaI*2, respectively. (C) A putative *lux*-box is present in the promoter region of the *rsaM*1 gene, which is homologous to characterized *lux*-box sequences in Proteobacteria.

**Figure S2.**
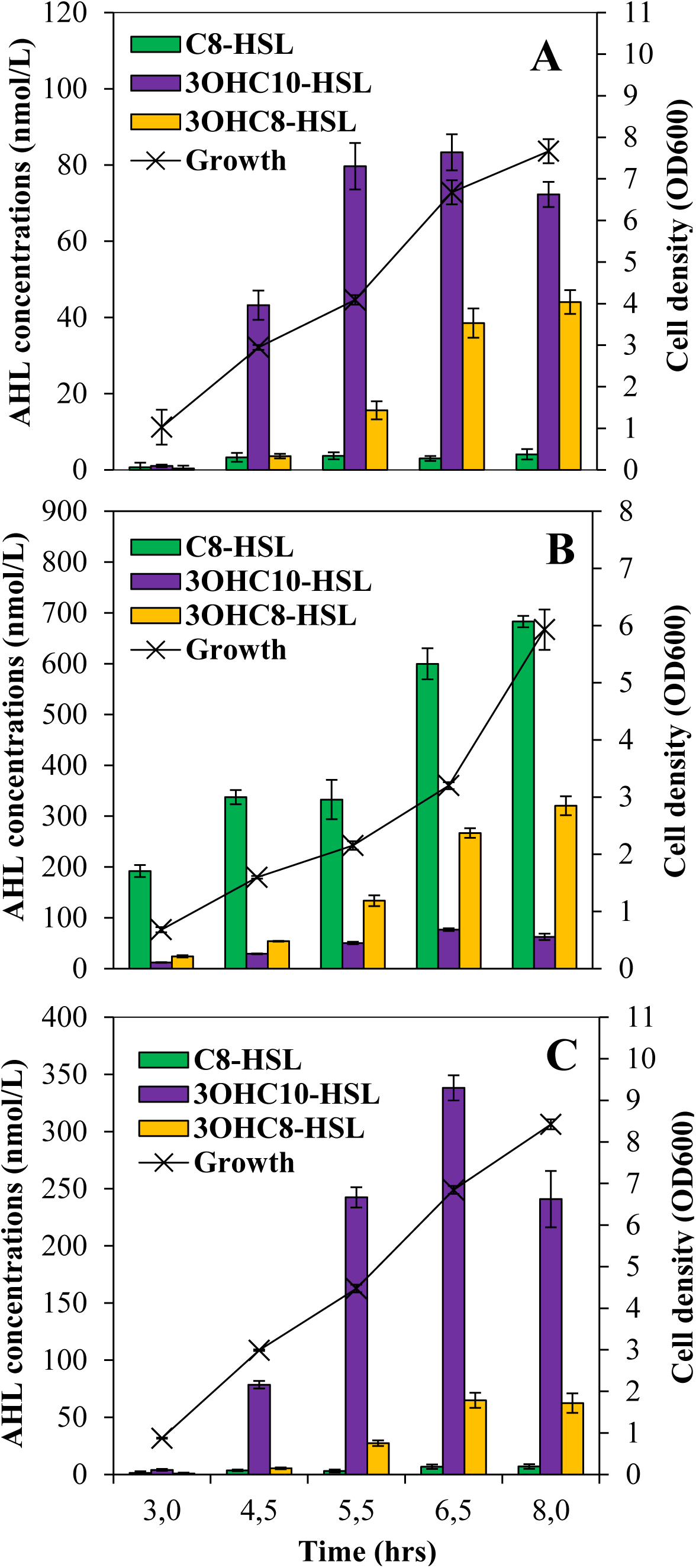
AHLs production profiles in the wild-type and the *rsaM***1- and *rsaM*2-mutant strains of *B. thailandensis* E264**. The biosynthesis of AHLs (bars) was monitored by LC-MS/MS at various times during growth (lines) in cultures of the (A) wild-type and the (B) *rsaM*1- and (C) *rsaM*2-mutant strains of *B. thailandensis* E264. The error bars represent the standard deviation of the average for three replicates.

**Figure S3.**
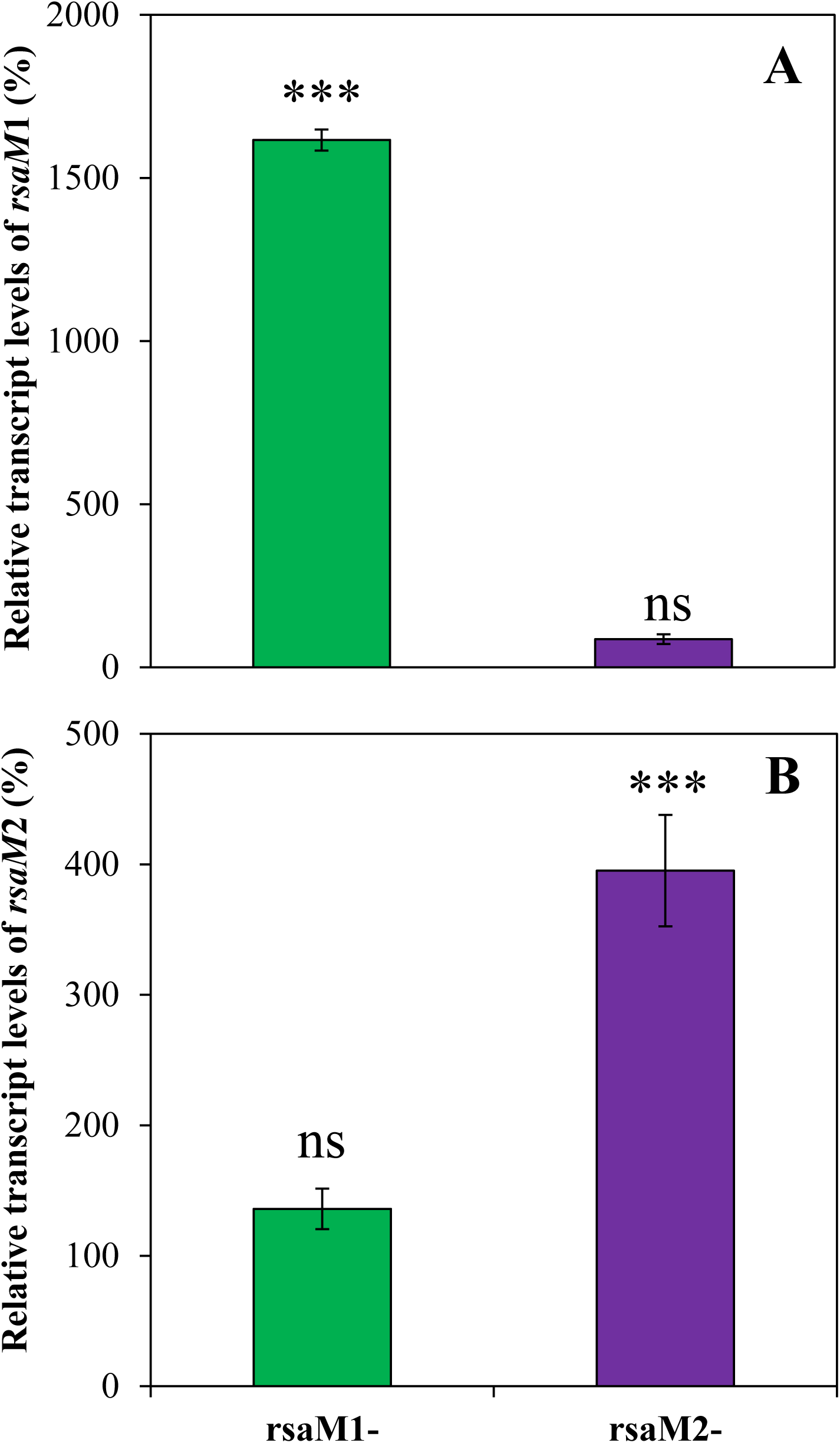
Expression of *rsaM***1 and *rsaM*2 is negatively auto-regulated**. The relative transcript levels of (A) *rsaM*1 and (B) *rsaM*2 from the *B. thailandensis* E264 wild-type and its *rsaM*1- and *rsaM*2-mutant strains were estimated by qRT-PCR experiments. The results are presented as relative quantification of genes transcription compared to the wild-type normalized to 100%. The error bars represent the standard deviation of the average for three replicates. ***, p < 0.001; ns, non-significant.

**Figure S4.**
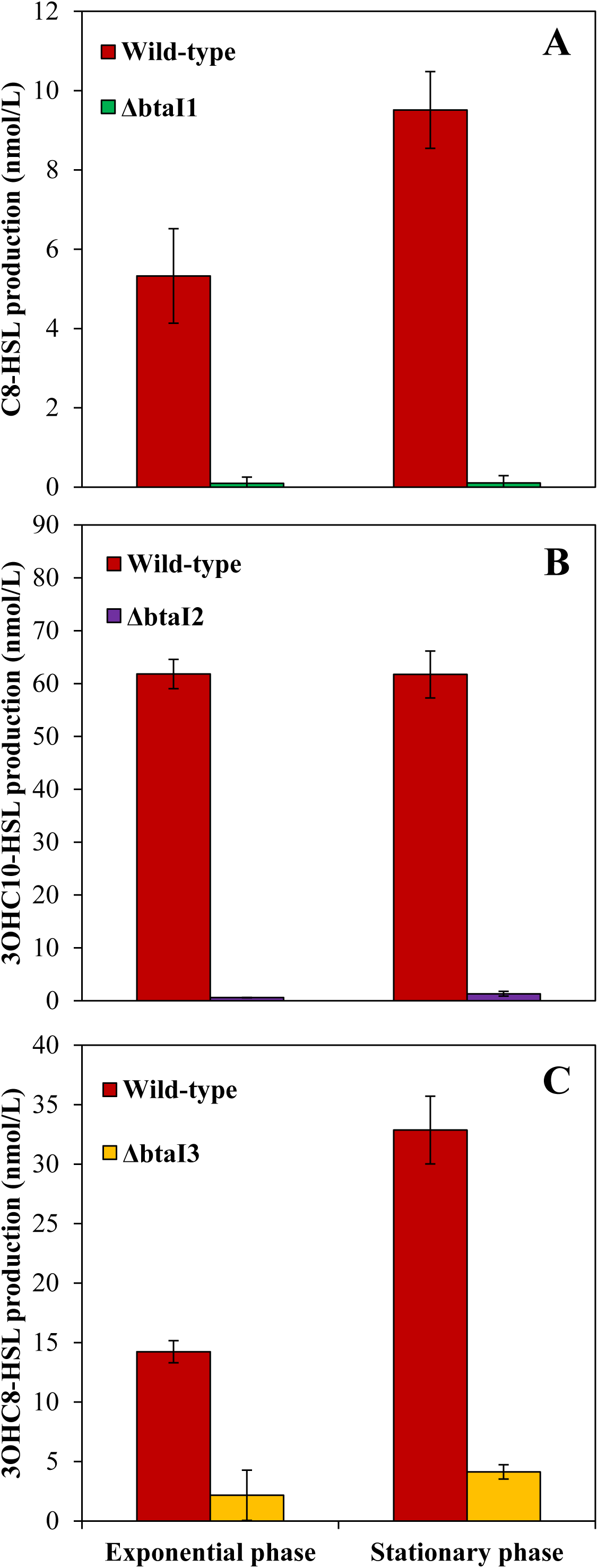
AHL biosynthesis in the wild-type and the Δ***btaI*1, Δ*btaI*2, and Δ*btaI*3 mutant strains of *B. thailandensis* E264**. The production of (A) C_8_-HSL, (B) 3OHC_10_-HSL, and (C) 3OHC_8_-HSL was quantified using LC-MS/MS during the exponential and stationary phases in cultures of the wild-type and of the ΔbtaI1, ΔbtaI2, and ΔbtaI3 mutant strains of *B. thailandensis* E264, respectively. The error bars represent the standard deviation of the average for three replicates.

**Figure S5.**
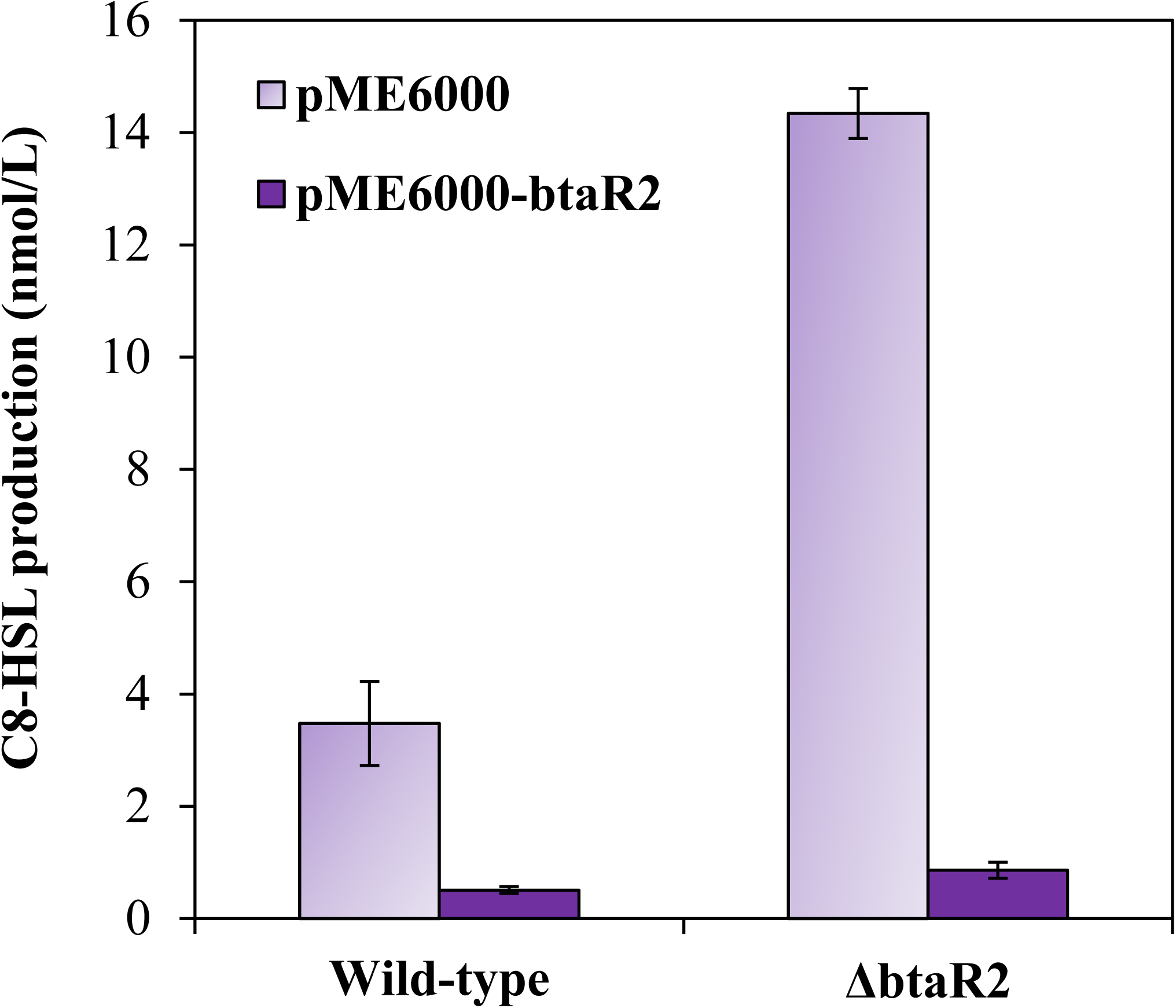
Complementation of C_8_-HSL production by BtaR2. The biosynthesis of C_8_-HSL was quantified using LC-MS/MS during the logarithmic growth in cultures of the wild-type and of the ΔbtaR2 mutant strains of *B. thailandensis* E264. The error bars represent the standard deviation of the average for three replicates.

**Figure S6.**
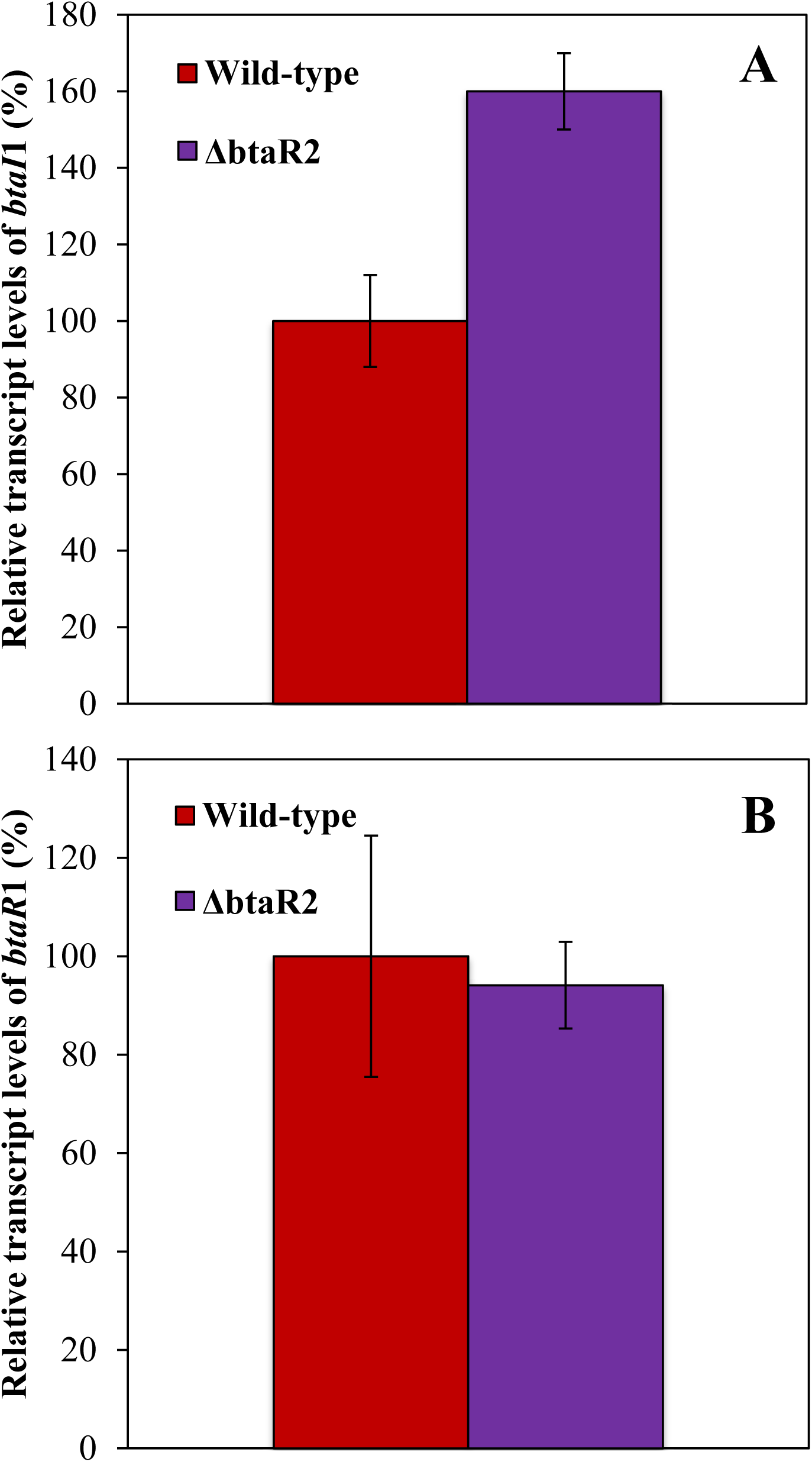
Expression of *btaI***1 is positively modulated by BtaR2**. The relative transcript levels of (A) *btaI*1 and (B) *btaR*1 from the *B. thailandensis* E264 wild-type and its Δ*btaR*2 mutant strains were estimated by qRT-PCR experiments during the logarithmic growth. The results are presented as relative quantification of genes transcription compared to the wild-type normalized to 100%. The error bars represent the standard deviation of the average for three replicates.

**Table S1.** Bacterial strains used in this study.

| Strains | Description | Reference |
| --- | --- | --- |
| *E. coli* |  |  |
| **χ7213** | *thr-1, leuB6, fhuA21, lacY1, glnV44, recA1, ΔasdA4, Δ(zhf-2::Tn10), thi-1, RP4-2-Tc :: Mu [λ pir]* | Lab collection |
| **DH5α** | F-, φ80dlacZΔM15, Δ(*lacZYA-argF*)U169, *deoR, recA1, endA1, hsdR17(rk-, mk+), phoA, supE44, λ-, thi-1, gyrA96, relA1* | Lab collection |
| *B. thailandensis* |  |  |
| **E264** | Wild-type | (33) |
| **JBT107** | E264 Δ*btaR1* | (16) |
| **JBT108** | E264 Δ*btaR2* | (16) |
| **JBT109** | E264 Δ*btaR3* | (16) |
| **JBT112** | E264 Δ*btaI1* Δ*btaI2* Δ*btaI3* | (16) |
| **BT03295** | E264 *rsaM1::ISlacZ-PrhaBo-Tp/FRT; Tp<sup>R</sup>* | (34) |
| **BT04218** | E264 *rsaM2::ISlacZ-PrhaBo-Tp/FRT; Tp<sup>R</sup> -* | (34) |
| **JBT101** | E264 Δ*btaI1* | (14) |
| **JBT102** | E264 Δ*btaI2* | (14) |
| **JBT103** | E264 Δ*btaI3* | (14) |
| **ED1020** | E264 (pME6000) | This study |
| **ED3312** | E264 (pME6000-*btaR2*) | This study |
| **ED1015** | E264 Δ*btaR2* (pME6000-*btaR2*) | This study |
| **ED3313** | E264 Δ*btaR2* (pME6000) | This study |
| **ED1015** | E264 Δ*btaR2* (pME6000-*btaR2*) | This study |
| **ED3226** | E264::*btaI1-lux* | (13) |
| **ED3306** | E264 *rsaM1*::*btaI1-lux* | This study |
| **ED3307** | E264 *rsaM2*::*btaI1-lux* | This study |
| **ED3227** | E264::*btaI2-lux* | (13) |
| **ED3308** | E264 *rsaM1*::*btaI2-lux* | This study |
| **ED3309** | E264 *rsaM2*::*btaI2-lux* | This study |
| **ED3228** | E264::*btaI3-lux* | (13) |
| **ED3310** | E264 *rsaM1*::*btaI3-lux* | This study |
| **ED3311** | E264 *rsaM2*::*btaI3-lux* | This study |

**Table S2.** Plasmids used in this study.

| Plasmids | Description | Source |
| --- | --- | --- |
| **pME6000** | Broad-host-range cloning vector; Tc<sup>R</sup> | (17) |
| **pMCG21** | *btaR2* inserted in *Bam*HI-*Hind*III sites in pME6000; Tc<sup>R</sup> | This study |
| **mini-CTX-*lux*** | Integration vector with promoterless *luxCDABE*; Tc<sup>R</sup> | (35) |
| **pSLG02** | *btaI1* promoter inserted in *Xho*I-*Bam*HI sites in mini-CTX-*lux*; Tc<sup>R</sup> | (13) |
| **pSLG03** | *btaI2* promoter inserted in *Xho*I-*Bam*HI sites in mini-CTX-*lux*; Tc<sup>R</sup> | (13) |
| **pSLG04** | *btaI3* promoter inserted in *Xho*I-*Bam*HI sites in mini-CTX-*lux*; Tc<sup>R</sup> | (13) |

**Table S3.**
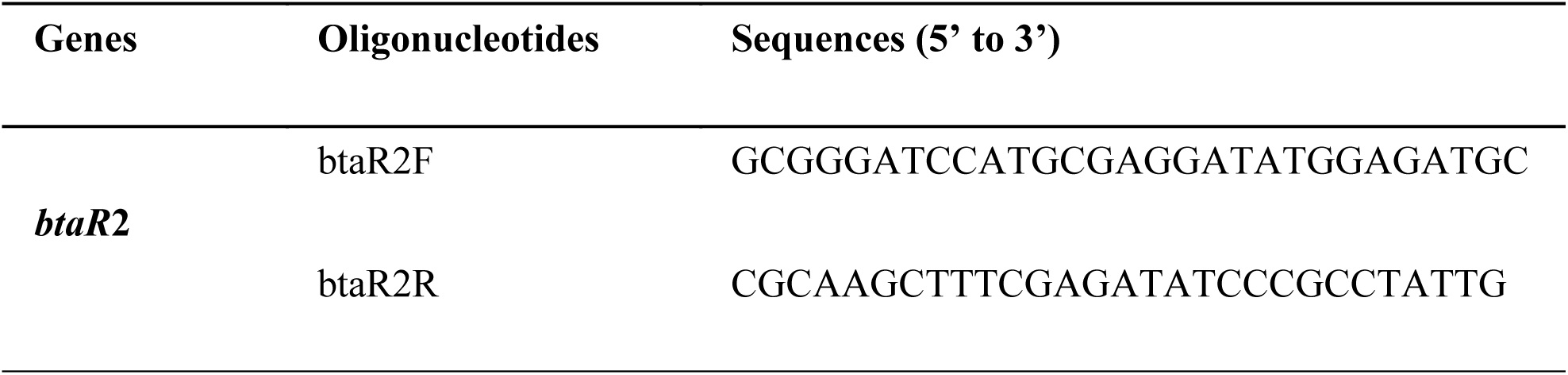
Primers used for PCR.

**Table S4.**
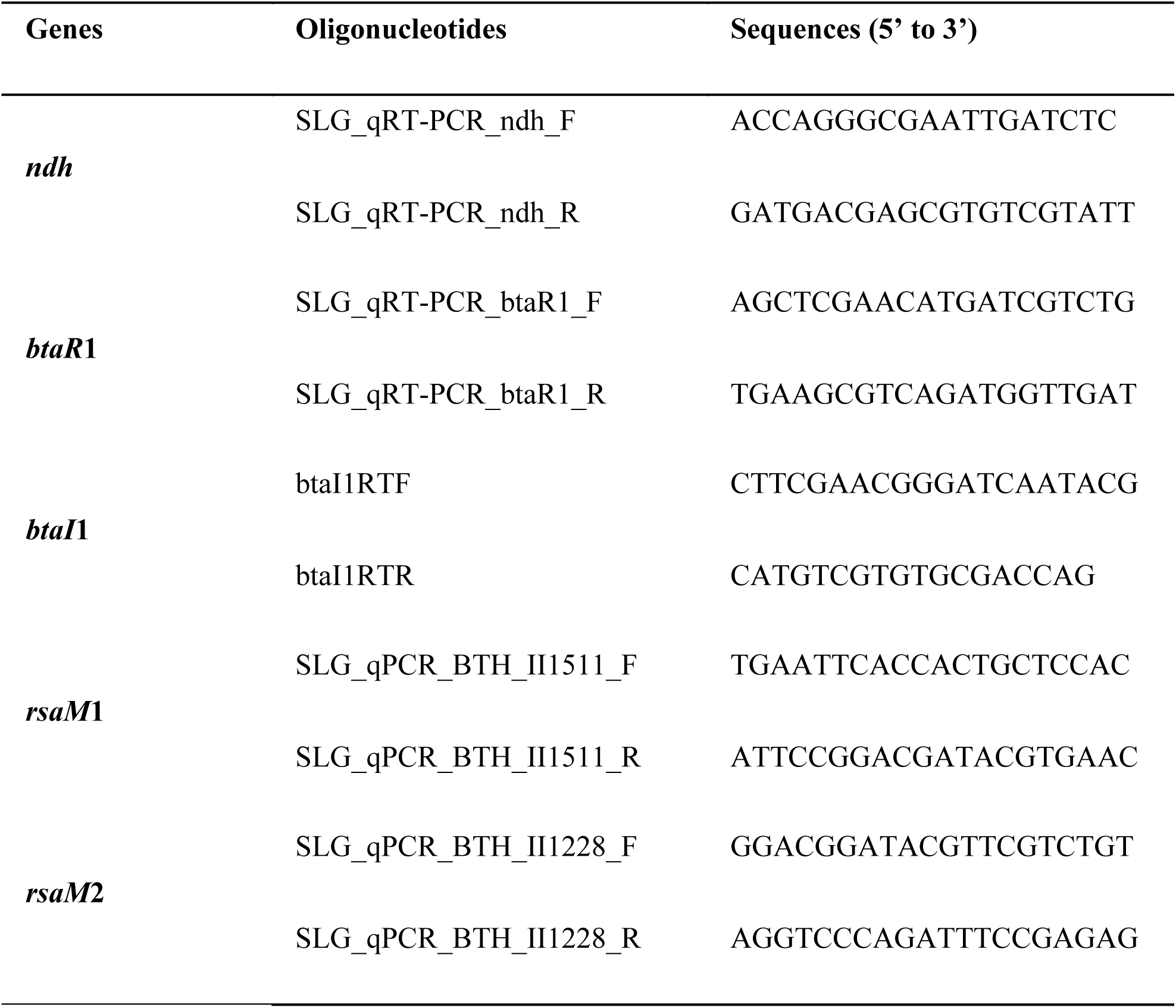
Primers used for qRT-PCR.

